# Placental mammals acquired the functional region and domain in NRK for regulating the CK2-PTEN-AKT pathway and placental cell proliferation

**DOI:** 10.1101/2021.09.22.461318

**Authors:** Beni Lestari, Satomi Naito, Akinori Endo, Hidenori Nishihara, Akira Kato, Erika Watanabe, Kimitoshi Denda, Masayuki Komada, Toshiaki Fukushima

## Abstract

The molecular evolution processes underlying the acquisition of the placenta in eutherian ancestors are not fully understood. Mouse NCK-interacting kinase (NIK)-related kinase (NRK) is expressed highly in the placenta and plays a role in preventing placental hyperplasia. Here, we show the molecular evolution of NRK, which confers its function for inhibiting placental cell proliferation. Comparative genome analysis identified NRK orthologues across vertebrates, which share the kinase and citron homology (CNH) domains. Evolutionary analysis revealed that NRK underwent extensive amino acid substitutions in the ancestor of placental mammals and has been since conserved. Biochemical analysis of mouse NRK revealed that the CNH domain binds to phospholipids, and a region in NRK binds to and inhibits casein kinase-2 (CK2), which we named the CK2-inhibitory region (CIR). Cell culture experiments suggest the following: (1) mouse NRK is localised at the plasma membrane via the CNH domain, where the CIR inhibits CK2. (2) This mitigates CK2-dependent phosphorylation and inhibition of PTEN, and (3) leads to the inhibition of AKT signalling and cell proliferation. *Nrk* deficiency increased phosphorylation levels of PTEN and AKT in mouse placenta, supporting our hypothesis. Unlike mouse NRK, chicken NRK did not bind to phospholipids and CK2, decrease phosphorylation of AKT, or inhibit cell proliferation. Both the CNH domain and CIR have evolved under purifying selection in placental mammals. Taken together, our study suggests that placental mammals acquired the phospholipid-binding CNH domain and CIR in NRK for regulating the CK2-PTEN-AKT pathway and placental cell proliferation.

## Introduction

The placenta is the defining organ of eutherians (placental mammals). The eutherian placenta is evolutionarily novel, is derived from the chorioallantoic membrane, and varies in morphology across species. It mediates the exchange of nutrients and gas between the mother and the foetus, regulates pregnancy as an endocrine tissue, and plays a role in suppressing the mother’s immune system (Imakawa and Nakagawa 2017). The development of the eutherian placenta is regulated by various extracellular or intracellular factors (Woods et al. 2018; Knofler et al. 2019). However, the molecular mechanisms by which they regulate placental development are not fully understood. Elucidating these molecular mechanisms will provide a basis for developing diagnostic and therapeutic methods for pregnancy complications, including foetal growth retardation. Moreover, it is also not fully understood how molecular evolution underlay the acquisition of the placenta in eutherian ancestors. Eutherian ancestors seem to have acquired several novel genes that code proteins involved in placental development, such as those derived from retrotransposons (Ono et al. 2006; Rawn and Cross 2008; Bolze et al. 2017; Imakawa and Nakagawa 2017). This has led to the increasing importance of research into the molecular evolution of proteins that regulate placental development.

The NCK-interacting kinase (NIK)-related kinase (NRK) is involved in mouse placental development. The mature mouse placenta comprises three histologically distinct layers: the labyrinth, spongiotrophoblast, and maternal decidual layers (Rossant and Cross 2001). Mouse *Nrk* is highly expressed in spongiotrophoblasts during late pregnancy, when the placenta weight reaches a plateau (Denda et al. 2011). Loss of *Nrk* leads to placental hyperplasia due to the hyperproliferation of the spongiotrophoblasts and difficulty during delivery (Denda et al. 2011). These observations suggest that mouse NRK plays a role in preventing hyperplasia of the placenta. Spongiotrophoblasts produce several hormones (e.g., placental lactogens) and regulate nutrient distribution between the foetus and placenta (Tunster et al. 2016). Mouse NRK may regulate the endocrine and nutritional environment in pregnancy by controlling spongiotrophoblast proliferation.

NRK is a member of the Germinal Center Kinase (GCK) family subgroup IV. This kinase family exhibits an *N*-terminal kinase domain, an uncharacterised middle region, and a *C*-terminal citron homology (CNH) domain (Delpire 2009). In *C. elegans* and flies, this family comprises only one member, namely, MIG-15 and MSN, respectively. They function as MAP kinase kinase kinase kinase (MAP4K), which is similar to the prototype Ste20 in yeast, and are involved in morphogenic events, including cell migration (Chapman et al. 2008; Plutoni et al. 2019). In mice and humans, the family comprises four kinases, namely, NIK, TRAF2- and NCK-interacting protein kinase (TNIK), misshapen-like kinase 1 (MINK1), and NRK (Delpire 2009). Experiments with mice and human cells have revealed that only NRK suppresses cell proliferation (Yu et al. 2014; Daulat et al. 2016; Masuda et al. 2016; Morioka et al. 2017; Naito et al. 2020). However, the evolutionary origin and molecular determinants of eutherian NRK function remain poorly understood.

Recently, we and the other research groups observed the ability of mouse NRK in suppressing AKT signalling, one of the major signalling pathways involved in the regulation of proliferation (Morioka et al. 2017; Naito et al. 2020). Phosphatidylinositol-3 kinase (PI3K), an upstream factor of the AKT signalling, is activated in response to growth factor stimuli and converts phosphatidylinositol 4,5-diphosphate (PIP_2_) to phosphatidylinositol 3,4,5-triphosphate (PIP_3_). AKT then recognises PIP_3_ and localises at the plasma membrane, where threonine 308 (T308) and serine 473 (S473) in AKT are phosphorylated by PDK1 and mTORC2, respectively. This induces the activation of AKT kinase, which phosphorylates various substrate proteins and ultimately promotes cell proliferation (Risso et al. 2015). Mouse NRK suppresses AKT phosphorylation (Morioka et al. 2017; Naito et al. 2020). However, the underlying mechanism remains elusive.

AKT signalling is regulated by various pathways, including the CK2-PTEN-AKT pathway. CK2 is a tetrameric protein complex consisting of two molecules of the catalytic subunit (α or α’ subunit) and two molecules of the regulatory subunit (β subunit). The CK2 complex may contain identical (two α or two α’) or non-identical (a pair of α and α’) catalytic subunits. CK2α and α’ exhibit overlapped functions. CK2 is ubiquitously expressed, is observed in various cellular compartments, and phosphorylates many substrate proteins (Litchfield 2003). PTEN, a lipid phosphatase that dephosphorylates the third phosphate group of PIP_3_, is one of the substrate proteins of CK2. CK2 phosphorylates multiple serine/threonine residues (S370, S380, T382, T383, and S385) near the C-terminus of PTEN. Unphosphorylated-PTEN demonstrates high phosphatase activity and is localised at the plasma membrane, thereby converting PIP_3_ to PIP_2_. In contrast, phosphorylated PTEN shows reduced phosphatase activity and is localised in the cytosol and the nucleus. Through these mechanisms, CK2 inhibits PTEN and increases PIP_3_ levels at the plasma membrane, thereby enhancing AKT signalling (Vazquez et al. 2000; Torres and Pulido 2001; Rahdar et al. 2009). However, the types of physiological situations in which AKT signalling is regulated via the CK2-PTEN-AKT pathway remains unclear.

This study aimed to reveal molecular evolution of the placental protein NRK, which confers its function for inhibiting placental cell proliferation.

## Materials and Methods

### Synteny analysis

We evaluated genomic data using Ensembl (https://www.ensembl.org) and the University of California, Santa Cruz (UCSC) genome browser (http://genome.ucsc.edu). We organised vertebrate *Nrk* and the neighbouring genes and identified syntenic genes by referring to the neighbouring genes located around *Nrk* in the mouse and chicken genome. We also identified syntenic genes around vertebrate *Nik*, *Tnik*, and *Mink1* genes by referring to the neighbouring genes located around *Nik*, *Tnik*, and *Mink1* genes in the human genome. Non-syntenic genes smaller than 10 kb in genomic size were omitted as such genes may have been mapped by mistake in gene prediction.

### Exon structure analysis

We retrieved the validated coding sequences of the human and mouse *Nrk* from the National Center for Biotechnology Information (NCBI; https://www.ncbi.nlm.nih.gov). The therian *Nrk* coding regions were identified from the genome sequences in **Supplementary Table 1** by referring to the human and mouse exon structures. RNA-seq data were obtained from NCBI SRA for platypus (SRX081892 and SRX122683), chicken (ERX697668 and ERX2593575), and zebrafish (DRX045951 and DRX045952); each species data were assembled using Bridger (Chang et al. 2015). The *Nrk* coding regions in the other vertebrates were identified from the genome sequences in **Supplementary Table 1** by referring to the above RNA-seq assemblies and the gene prediction data in NCBI (platypus: XM_029067556.1, *Xenopus*: XM_031891524.1, spotted gar: XM_006632839.2, and reedfish: XM_028815688.1) and Ensembl (painted turtle: ENSCPBT00000034663.1, coelacanth: ENSLACT00000019211.1, and zebrafish: ENSDARG00000098680). We searched the origin of the inserted sequences in the transposable element collection (Repbase repeat library) using RepeatMasker (http://www.repeatmasker.org).

### Protein evolutionary rate analysis

The NRK amino acid sequences from 24 species were aligned for each exon using MAFFT version 7 (Katoh and Standley 2013) with the linsi option followed by partial adjustments with MUSCLE implemented in MEGA (Tamura et al. 2021). We reconstructed the ancestral amino acid sequences by using FastML (Ashkenazy et al. 2012) with the JTT+G model and ML-based indel reconstruction under the assumption of known species tree in TimeTree (accessed on 5 August 2020; Kumar et al. 2017). The number of amino acid substitutions was counted for each branch by comparing the estimated ancestral sequences of the two corresponding nodes, and the substitution rates were calculated based on the divergence times available in TimeTree. The estimated indel information was used only for visualisation and ignored in the substitution rate calculations. Neighbour-joining tree for the amino acid sequences of the NRK domains were estimated under the JTT+G model with the pairwise deletion option.

### Gene expression analysis

We collected gene expression data of mouse *Nik*, *Tnik*, *Mink1*, *Nrk,* and human *NRK* in various tissues analysed by the cap analysis of gene expression (CAGE) in RIKEN FANTOM5 project from the public database RefEx (Reference Expression dataset; http://refex.dbcls.jp; Ono et al. 2017) and Expression Atlas (https://www.ebi.ac.uk/gxa/home). We also collected gene expression data of sheep (Jiang et al. 2014), chicken (Merkin et al. 2012), and *Xenopus Nrk* (Barbosa-Morais et al. 2012) in various tissues from the Expression Atlas. The data were then visualised using an online tool, Heatmapper (http://heatmapper.ca/expression/).

### dN/dS calculation

The ratio of non-synonymous to synonymous substitution (dN/dS) was calculated to estimate the selective constraint of the *Nrk* sequences in 15 eutherian mammals (human, orangutan, tarsier, mouse, squirrel, dog, cat, horse, microbat, minke whale, sheep, cow, shrew, elephant, and tenrec) under the species tree topology. This calculation was performed based on the Bayes Empirical Bayes method with the M8 model by using CODEML in the Phylogenetic Analysis by Maximum Likelihood (PAML) 4.9j package (Yang 2007).

### Antibodies and reagents

The following primary antibodies were used in this study: anti-FLAG M2 antibody (clone M2, Sigma-Aldrich, St Louis, MO, USA), anti-myc antibody (clone 9E10, Santa Cruz Biotechnology, Dallas, TX, USA), HRP-conjugated anti-GST antibody (#RPN1236, Cytiva, Marlborough, MA, USA), anti-NRK rabbit serum (Denda et al. 2011), anti-PARP antibody (clone C-2-10, Calbiochem, San Diego, CA, USA), anti-AKT antibody (clone C67E7, Cell Signalling Technology (CST), Danvers, MA, USA), anti-phospho AKT (T308) antibody (clone D25E6, CST), anti-phospho AKT(S473) antibody (clone D9E, CST), anti-p42/44 MAPK antibody (#9102, CST), anti-phospho p42/44 MAPK (T202/T204) antibody (clone E10, CST), anti-CK2α antibody (clone 1AD9, Santa Cruz Biotechnology), anti-CK2α’ antibody (A300-199A, Bethyl Laboratories, Montgomery, TX, USA), anti-CK2β antibody (A301-984A, Bethyl Laboratories), anti-phospho CK2 substrate antibody (#8738, CST), anti-PTEN antibody (clone 138G6, CST), anti-phospho PTEN (S380) antibody (clone H-3, Santa Cruz Biotechnology), anti-α-Tubulin antibody (#013-25033, Wako Pure Chemical, Osaka, Japan), anti-phosphatidylinositol 4,5-bisphosphate (PIP_2_) antibody (clone 2C11, Santa Cruz Biotechnology), and anti-phosphatidylinositol 3,4,5-triphosphate PIP_3_ (#Z-P345, Echelon Biosciences, Salt Lake City, UT, USA). For immunoprecipitation, anti-FLAG M2 affinity gel (Sigma-Aldrich) was used. Peroxidase-labelled anti-mouse IgG antibody and anti-rabbit IgG antibody (GE Healthcare, Chicago, IL, USA) were used as secondary antibodies for immunoblotting. Alexa Fluor 488- or 594-conjugated anti-mouse IgG antibody, anti-mouse IgM antibody, and anti-rabbit IgG antibody (Thermo Fisher Scientific, Waltham, MA, USA) were used as secondary antibodies for immunostaining. Cells were treated with the following reagents: doxycycline (dox; Sigma-Aldrich), epidermal growth factor (EGF; PeproTech, Rocky Hill, NJ, USA), cycloheximide (Nacalai Tesque, Kyoto, Japan), tumour necrosis factor-α (TNFα; Suntory, Tokyo, Japan), and a PTEN inhibitor VO-OHpic (BioVision, Milpitas, CA, USA).

### Plasmids

Mouse *Nrk* cDNA was pre-prepared by our group (Nakano et al. 2000). Chicken *Nrk* cDNA (NM_001031126.2) was amplified using polymerase chain reaction (PCR) from the chicken embryo cDNA library kindly donated by Dr. Mikiko Tanaka (Tokyo Institute of Technology, Yokohama, Japan). Mouse *Nik* (XM_006496042.3), *Tnik* (NM_026910.1), *Mink1* (NM_016713.2), *CK2*α*’* (see below), *CK2*β (NM_001303445.1), and *PTEN* (NM_008960.2) cDNAs were amplified by using PCR from mouse embryo and brain cDNA libraries. The mouse *CK2*α*’* cDNA sequence is identical to NM_009974.3, except for the 84 bp deletion in the predicted coding region, which is thought to be an isoform because other rodent *CK2*α*’* cDNA sequences also lack this 84 bp sequence. The following vectors were used in this study: mammalian expression vectors, pFLAG-CMV2 (Sigma-Aldrich) and pCMV5 (kindly donated by Dr. Jun Nakae, Keio University, Tokyo, Japan); dox-induced mammalian expression vector, pcDNA5/FRT/TO (Invitrogen, Carlsbad, CA, USA); bacterial expression vectors, pGEX-4T1 and pGEX-6P2 (GE Healthcare). N-terminal FLAG-tagged mouse NRK (mNRK), mNRK CNH (aa 1133-1455), mNRK Δ CNH (aa 1-1137), mNRK Δ CNH-CaaX (mNRK aa 1-1137 fused with HRAS CaaX motif (GCMSCKCVLS) at the C-terminus), other mNRK truncated mutants, mouse NIK (mNIK), mNIK CNH (aa 1011-1330), mouse TNIK (mTNIK), mTNIK CNH (1042-1361), mouse MINK1 (mMINK1), mMINK1 CNH (982-1301), chicken NRK (cNRK), and cNRK CNH (aa 918-1237) were expressed using pFLAG-CMV2 or pcDNA5/FRT/TO. N-terminal myc-tagged mouse CK2α’ and N-terminal myc-tagged mouse CK2β were expressed using pCMV5. GST-tagged mNRK (aa 565-868) was expressed using pGEX-6P2 (Cytiva). GST-tagged NRK (aa 565-831), GST-tagged PTEN (aa 190-403), GST-tagged mNRK CNH (aa 1133-1455), and GST-tagged cNRK CNH (aa 918-1237) were expressed using pGEX-4T1 (Cytiva).

### Cell culture and transfection

HEK293 cells that express mNRK in a dox-dependent manner were developed using Flp-In T-REx 293 cells (Invitrogen), according to the manufacturer’s instructions. They were cultured in Dulbecco’s modified Eagle’s medium (DMEM) supplemented with 10% foetal bovine serum (FBS), 100 U/mL penicillin, 100 μg/mL streptomycin, 7.5 μ /mL blasticidin (Wako), and 16 μg/mL hygromycin (Invitrogen). HEK293, HEK293T, and HeLa cells were cultured in DMEM supplemented with 10% FBS, penicillin, and streptomycin. DNA transfection was performed using polyethylenimine (PEI, linear; molecular weight, 25,000; Polyscience, Warrington, PA, USA).

### MTT assay

HEK293 cells expressing NRK in a dox-dependent manner were seeded in collagen-coated 96-well plates at a density of 2 × 10^3^/well and cultured for 24 h. The cells were treated with dox (1 μg/mL) and incubated for 48 h. A total of 10 µL of cell count reagent SF (#07553-15, Nacalai Tesque) was added, and cells were incubated for an additional 2 h. The absorbance was then measured at 450 nm using microplate readers (Varioskan^TM^ LUX, Thermo Fisher Scientific).

### Immunoprecipitation and immunoblotting

Cells were lysed with ice-cold lysis buffer (20 mM Tris-HCl, pH 7.4, 100 mM NaCl, 50 mM NaF, 0.5% Nonidet P-40, 1 mM EDTA, 1 mM phenylmethylsulfonyl fluoride, 1 µg/mL aprotinin, 1 µg/mL leupeptin, and 1 µg/mL pepstatin A). For the detection of phosphorylation signals, 1 mM Na_3_VO_4_ was added to the lysis buffer. The lysates were then centrifuged at 15,000 rpm for 15 min. The supernatants were subjected to immunoprecipitation and immunoblotting according to standard protocols. Immunoblotting signals were detected using the ECL prime western blotting detection reagents (GE Healthcare) and the ImageQuant LAS 4000 mini-imager (GE Healthcare). Densitometric analyses were performed using the ImageJ program (version 1.52).

### Immunostaining

Immunostaining analyses were performed according to standard procedures. Briefly, HeLa cells were cultured on glass coverslips and transfected with PEI. After 48 h of transfection, cells were fixed with 4% paraformaldehyde (PFA) in phosphate buffer saline (PBS) for 10 min at 20–25 °C, permeabilised with 0.2% Triton X-100 in PBS, and blocked with 5% FBS in PBS. The samples were then incubated with primary antibodies for 16 h at 4 °C, followed by secondary antibodies for 1 h at 20–25 °C. DAPI (1 μg/mL, Nacalai Tesque) was added at the end for nuclear staining before mounting. As an exception, immunostaining of PIP_2_ and PIP_3_ was performed according to the protocol provided by Echelon Biosciences. Briefly, the transfected HeLa cells were fixed with 4% PFA and permeabilised with 0.01% digitonin for 20 min each at 20–25 °C. After blocking with 10% goat serum in TBS for 30 min at 37 °C, cells were then incubated with anti-PIP_2_ or anti-PIP_2_ antibodies followed by secondary antibodies for 1 h each at 37 °C. DAPI was added at the end for nuclear staining before mounting. Fluorescence images were acquired using a laser-scanning confocal microscope (LSM 780, Carl Zeiss, Oberkochen, Germany).

### Lipid-protein overlay assay

GST-tagged CNH domains of mNRK and cNRK (GST-CNH) were expressed in *E. coli* strain Rosetta (DE3)pLysS and purified using Glutathione-Sepharose 4B beads (GE Healthcare). Membrane lipid strips (Echelon Biosciences) were blocked with blocking buffer containing PBS, 3% bovine serum albumin (BSA), and 0.1% Tween 20 for 1 h at 20–25 °C. Membranes were then incubated with 3 µg/mL of GST-CNH in blocking buffer for 1 h at 20–25 °C followed by washing thrice for 10 min each in a washing buffer containing PBS and 0.1% Tween 20. Subsequently, the membranes were incubated with HRP-conjugated antibody (Cytiva) for 1 h at 20–25 °C, washed thrice for 10 min each with washing buffer, and subjected to chemiluminescence detection using the ECL prime reagents and the ImageQuant LAS 4000 mini-imager.

### *In vitro* CK2 kinase assay

A purified CK2 complex was purchased (#P6010S, New England BioLabs, Beverly, MA, USA). GST-mNRK^565-868^, GST-mNRK^565-831^, and GST-PTEN^190-403^ fusion proteins were expressed in *E. coli* strain Rosetta (DE3)pLysS, adsorbed to Glutathione-Sepharose 4B beads, and eluted with reduced glutathione. Non-tagged mNRK^565-868^ was eluted with PreScission Protease (Cytiva) according to the manufacturer’s recommended protocol. SDS-PAGE and CBB staining confirmed the purity of proteins used in this study. For the *in vitro* kinase assay, CK2 (20 units) and the substrate GST-PTEN^190-403^ (1 µg) were mixed with recombinant mNRK proteins (0.5 µg) or bovine serum albumin (BSA, 0.25 µg) in a kinase assay buffer (20 mM Tris-HCl, pH 7.5, 5 mM MgCl_2_, 150 mM KCl, 0.5 mM DTT, 0.3 µM ATP) and incubated at 30 °C for 30 min. Reactions were stopped by the addition of SDS-PAGE sample buffer and boiling for 5 min. Samples were subjected to SDS-PAGE and immunoblotting.

To test whether mNRK^565-868^ may compete with PTEN as a substrate for CK2, mNRK^565-868^ (0.25 µg) or BSA (0.25 µg) was mixed with CK2 (10 units; denoted as CK2^1^) in the kinase assay buffer and preincubated at 30 °C for 16 h to prepare phosphorylated mNRK^565-868^ and the negative control. In some samples, the reactions were stopped just after preincubation. Other samples were mixed with CK2 (10 units; denoted as CK2^2^), GST-PTEN^190-403^ (1 µg), and ATP (0.3 µM) and incubated at 30 °C for 30 min. The other samples were incubated similarly except for the absence of CK2^2^. The reactions were stopped, and samples were analysed as described above.

### Animal experiment

Female heterozygous mice (X^-^X^+^) were mated with male null mice (X^-^Y). The placentas at 18.5 dpc were dissected, genotyped, and divided into two groups (WT, placentas of male foetuses [X^+^Y] and female foetuses [X^-^X^+^]; KO, placentas of male foetuses [X^-^Y] and female foetuses [X^-^X^-^]). Note that the paternally derived X chromosome is preferentially inactivated in the extraembryonic tissues of mice (Takagi and Sasaki 1975). Placentas were dissected at 18.5 days post-coitum (dpc) from pregnant female mice and cut into small pieces. Placental proteins were then lysed and sonicated in lysis buffer containing 20 mM Tris-HCl pH 7.4, 100 mM NaCl, 50 mM NaF, 1% Nonidet P-40, 0.1% sodium dodecyl sulphate, 1 mM EDTA, 1 mM phenylmethylsulfonyl fluoride, 1 µg/mL aprotinin, 1 µg/mL leupeptin, and 1 µg/mL pepstatin A. The lysates were subjected to immunoblotting. All animal experiments were approved and conducted in compliance with the Institutional Animal Care and Research Advisory Committee of the Tokyo Institute of Technology guidelines.

## Statistical analysis

The number of independent experiments is indicated in figure legends. All data are presented as mean ± standard deviation (SD) of independent experiments. Graphs and statistical analyses were processed using GraphPad Prism 9. Asterisks on figures indicate values that were statistically different (*p≤0.05; **p≤0.01; ***p≤0.001; ****p≤ 0.0001).

## Results

### *Nrk* orthologues were identified across vertebrates

Our synteny analysis showed that the *Nrk* gene was widely present in vertebrates, including eutherians, marsupials, birds, reptiles, amphibians, and fish (**Fig. 1A**). We also found that the genes of other GCK IV family members (*Nik*, *Tnik*, and *Mink1*) were widely present in mammals, birds, reptiles, amphibians, and fish (**Supplementary Fig. 1A** and **Supplementary Table 2**). Notably, we did not find the *Mink1* gene in chicken and turtle genomes. These results indicate that each gene of the GCK IV family members was present in the common ancestor of vertebrates.

**Fig. 1.**
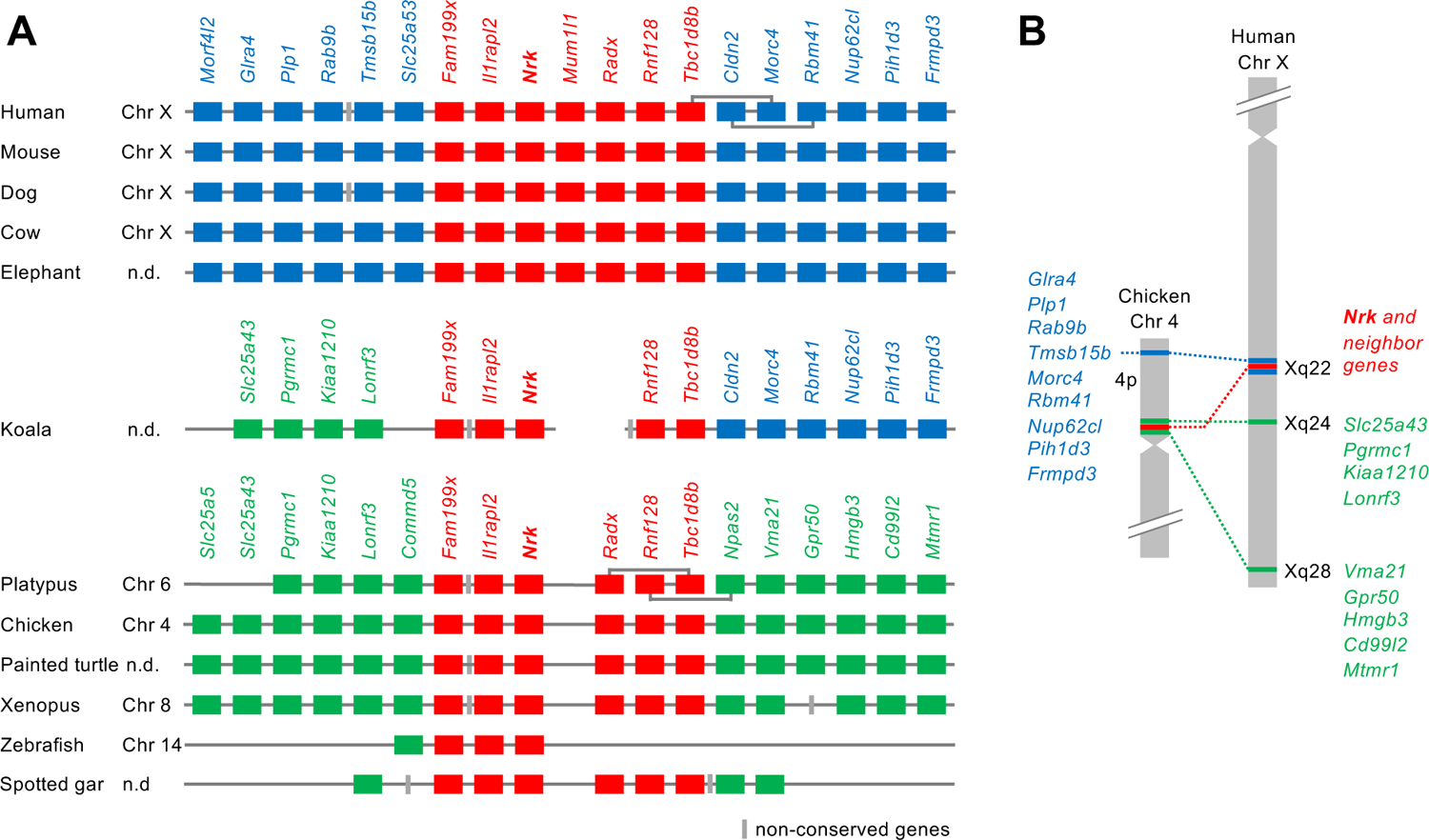
Synteny analysis of the *Nrk* gene. **(A)** Synteny conservation of the region surrounding the *Nrk* gene. Red, blue, and green boxes indicate syntenic genes across vertebrates, eutherians, and those observed only in platypus and non-mammals, respectively. Gray boxes indicate non-conserved genes. **(B)** Loci of *Nrk* and syntenic genes in chicken and human chromosomes.

#### Genomic region containing *Nrk* was relocated during early mammalian evolution

We compared gene loci and synteny of vertebrate *Nrk* orthologues (**Fig. 1A**). Human, mouse, dog, and cow *Nrk* genes were located on the X chromosome. In platypus and chicken, *Nrk* genes were located on chromosomes 6 and 4, which contain regions orthologous to the eutherian X chromosome (Veyrunes et al. 2008). Synteny analysis of *Nrk* and the neighbouring genes showed three different groups of syntenic genes. Red-coloured genes (*Nrk* and some neighbouring genes) retain their synteny from *Xenopus* to humans. In eutherians, blue-coloured genes upstream and downstream of red genes were syntenic. However, in platypus and non-mammals, a different group of genes (green genes) showed synteny. Incomplete koala genome data showed that the genomic region on one side of the *Nrk* gene was orthologous to the regions of platypus and non-mammals, and the genomic region on the other side was orthologous to certain regions of eutherians. Hence, the genomic region in the koala was maintained in a transitional state. The human Xq is highly homologous to the 20 Mb region of chicken 4p (Ross et al. 2005). We compared the loci of *Nrk* and neighbouring genes in chicken and human chromosomes (**Fig. 1B**). In chicken, the red and green genes were located near the centromere of 4p, and the blue genes were located near the end of 4p. In humans, the red and blue genes were located at Xq22. The green genes on the left side of *Nrk* in Fig. 1A were located at Xq24, whereas the green genes on the right side were located at Xq28.

### *Nrk* gene underwent rapid molecular evolution during early mammalian evolution

Next, we identified coding sequences of vertebrate *Nrk* genes in the orthologous loci (**Supplementary Table 1**) based on several public data of validated mRNA sequences, gene predictions, and RNA-seq assemblies (see Materials and Methods). We compared the exon structures of vertebrate *Nrk* orthologues, and classified them into three types (eutherian, marsupial, and platypus and non-mammalian; **Supplementary Fig. 2A**). The human *Nrk* gene consisted of 29 exons, all of which were coding exons. Almost all eutherian *Nrk* genes demonstrated orthologues for the aforementioned 29 exons. The therian exon 13 was longer than the orthologous exon of platypus and non-mammals. The orthologous sequences of the eutherian exons 4, 12, and, 25 were not found in non-eutherians. The eutherian exons 18 and 19 were longer than those of the non-eutherians. These changes led to the insertions of amino acid sequences that were found only in the eutherian NRK protein. We searched the origin of the inserted sequences in the transposable element database, but no similar element was found.

Using the *Nrk* coding sequences, we estimated the evolutionary rate of the amino acid sequences based on the divergence times of the species in TimeTree (Kumar et al. 2017; **Fig. 2**, left panel). We also analysed the distribution of amino acid changes in the NRK sequences occurring in each branch from the tetrapod ancestor to humans (**Fig. 2**, right panel; black, no change; gray, insertion; red, substitution). The amino acid evolutionary rate in the eutherian ancestor was 10.8 × 10^-9^ substitutions/site/year, much higher than those in the therian ancestor (2.6 × 10^-9^ substitutions/site/year) and the mammalian ancestor (0.012 × 10^-9^ substitutions/site/year). The results demonstrate an accelerated evolutionary rate in the eutherian ancestor, approximately 100–180 million years ago (Mya). During this period, the NRK sequence was lengthened owing to certain insertions as shown in the exon structure analysis. The protein sequence underwent many substitutions in several regions, even including the kinase and CNH domains. The evolutionary rates after the most recent common ancestor of eutherians were low (1.2 × 10^-9^ substitutions/site/year on average of the terminal branch), suggesting the sequence and functional conservation of *Nrk* among eutherians.

**Fig. 2.**
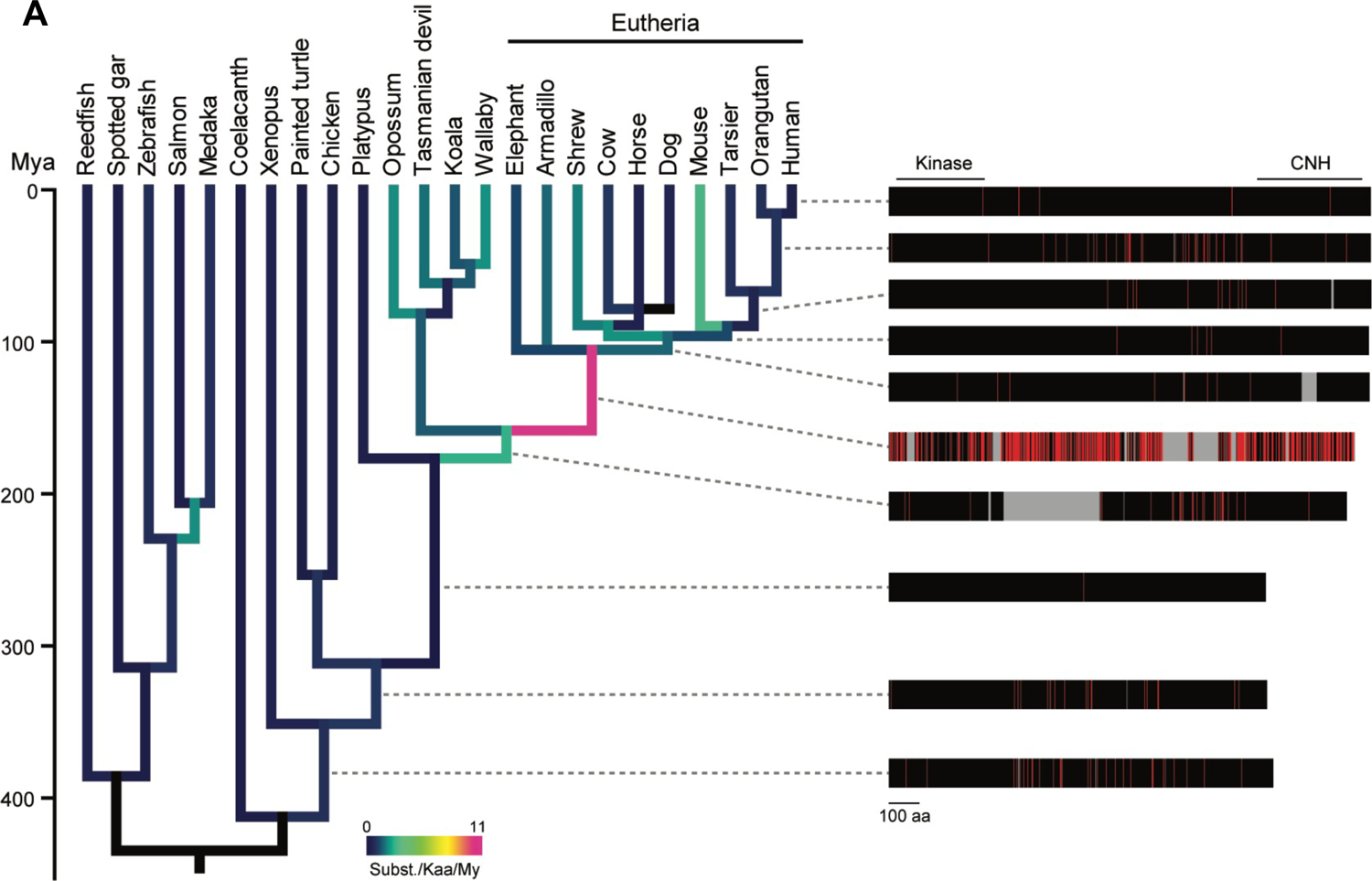
Phylogenetic analysis of the amino acid sequences of vertebrate NRK Amino acid substitutions of NRK along with the vertebrate phylogeny. The amino acid substitutions in each branch were estimated based on the reconstructed ancestral sequences of NRK proteins. The divergence times were retrieved from TimeTree. The colours of the branches indicate the average number of substitutions per 1,000 amino acid sequences over a million years. The right panel shows the changes in amino acid lengths and sequences of ancestral proteins of human NRK. The positions of the kinase and CNH domains of the human NRK are indicated. Sites of amino acid substitutions, unchanged sites, and insertions in each branch are shown in red, black, and gray, respectively. My, million years.

We found that vertebrate NRK protein shares a similar domain architecture: the *N*-terminal kinase domain and the *C*-terminal CNH domain. The neighbour-joining (NJ) trees for the amino acid sequences of these domains suggest that both domains of eutherians were distinct from those of non-eutherians (**Supplementary Fig. 2B, C**), consistent with the result of the evolutionary rate analysis.

### *Nrk* gene became to be expressed preferentially in the placenta during early mammalian evolution

Using a public database, we compared the tissue expression patterns of the mouse *Nrk* (*mNrk*) gene and the other GCK IV family members (*Nik*, *Tnik*, *and Mink1*). Mouse *Nrk* was restrictedly expressed in the placenta, whereas other genes were ubiquitously expressed (**Supplementary Fig. 3A**). We also compared the tissue expression pattern of *Nrk* orthologues in different species. The data were limited, but those of human, sheep, chicken, and *Xenopus* were available. Human and sheep *Nrk* genes were restrictedly expressed in the placenta. Chicken and *Xenopus Nrk* genes were highly expressed in other tissues (**Supplementary Fig. 3B**). These results suggest a eutherian-specific pattern of *Nrk* gene expression.

### Mouse NRK suppressed cell proliferation and AKT signalling, for which the middle region of NRK and the CNH domain are required

Next, we analysed the molecular function of mouse NRK (mNRK) and compared it with the functions of NRK orthologues and the other GCK IV family members. To the best of our knowledge, no monotypic cell line is known in which endogenous NRK protein can be detected. Using the HEK293 Flp-In T-REx cell system, we generated cells that expressed mNRK or green fluorescence protein (GFP) in a dox-dependent manner (Naito et al. 2020). Compared with the control cells, dox-treated cells showed decreased cell numbers (**Fig. 3A**) and inhibited AKT phosphorylation without affecting MAPK phosphorylation (**Fig. 3B**). Immunostaining analysis confirmed that mNRK expression inhibited AKT phosphorylation (**Fig. 3C**). mNRK expression did not induce cleavage of PARP1 (**Supplementary Fig. 4**), indicating that it did not mediate cellular apoptosis. We then examined the effects of the other GCK IV family members (mouse NIK, TNIK, and MINK1) and chicken NRK (cNRK) on AKT phosphorylation. The expression of these proteins did not decrease AKT phosphorylation (**Fig. 3D**). The protein levels were varied, probably owing to differences in protein stabilities under our experimental conditions. These results demonstrate the unique inhibitory function of mNRK in mediating AKT signalling.

**Fig. 3.**
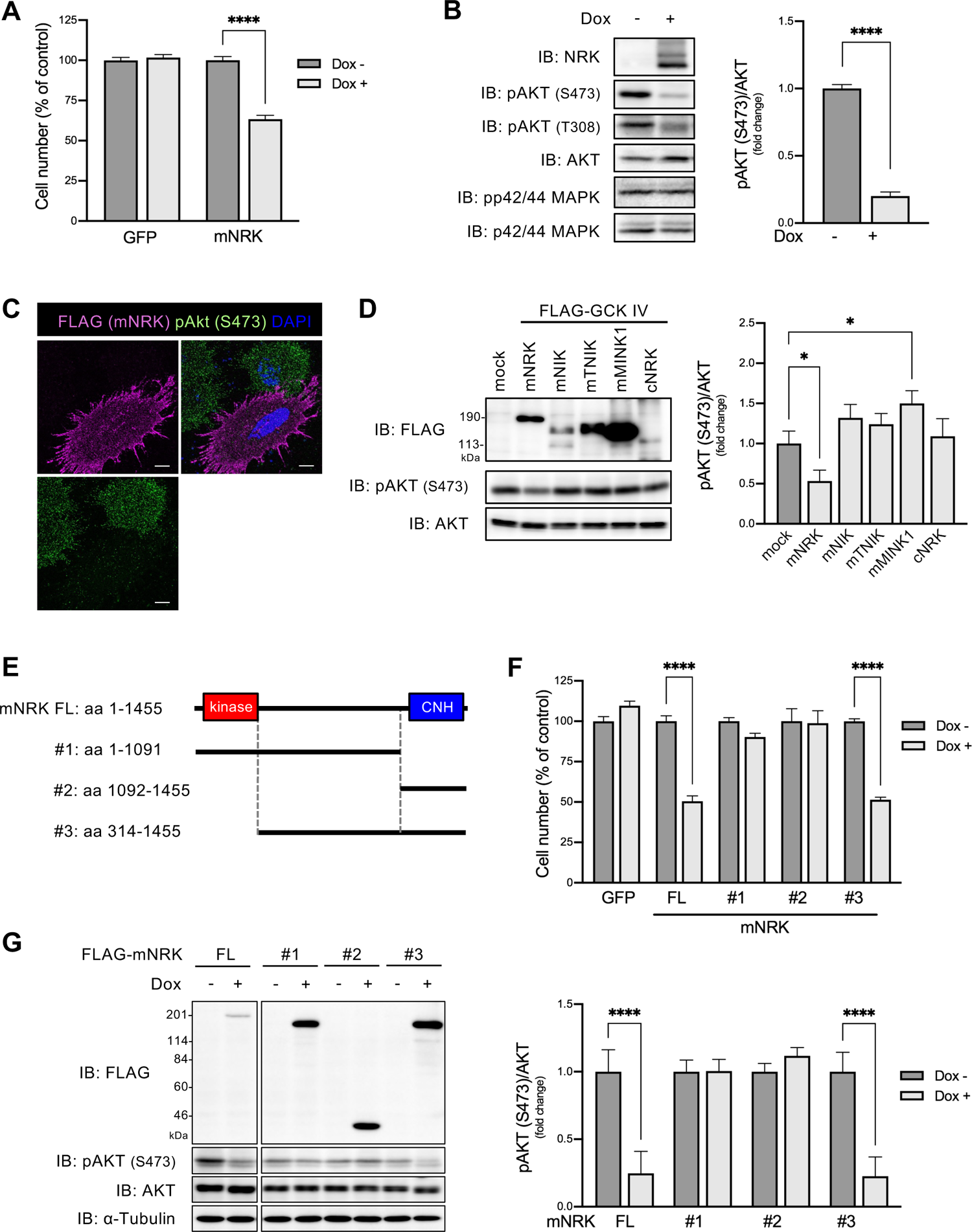
Effects of mNRK on cell proliferation and AKT signalling and the corresponding responsible regions of mNRK. **(A)** Effects of mNRK on cell proliferation. We generated Flp-In TREx 293 cell lines that expressed mNRK and GFP in doxycycline (dox)-dependent manner. The cells were treated with dox (1 μg/mL) for 2 days and then subjected to MTT assay. The graph shows the mean ± standard deviation (SD) of three independent experiments. ****, p ≤ 0.0001 (unpaired two-tailed Student’s t-test). **(B, C)** Effects of mNRK on AKT signalling. **B**) Cells prepared by a method similar to that shown in **A** were subjected to immunoblotting. Densitometric analyses were performed, and phosphorylation levels of AKT (S473) were normalised to protein levels of AKT. The graph shows the mean ± standard deviation (SD) of three independent experiments. ****, p ≤ 0.0001 (unpaired two-tailed Student’s t-test). **C)** HeLa cells expressing FLAG-tagged NRK were stimulated with EGF (100 ng/mL) for 10 min and subjected to cell staining with anti-FLAG antibody (magenta), anti-phospho AKT (S473) antibody (green), and DAPI (blue). Scale bar, 10 µm. **(D)** Effects of the other GCK IV family members (mNIK, mTNIK, mMINK1) and cNRK on AKT signalling. HEK293 cells were transfected with plasmids encoding mNRK, mNIK, mTNIK, mMINK1, and cNRK. The lysates were subjected to immunoblotting, followed by densitometric analyses. The graph shows the mean ± standard deviation (SD) of three independent experiments. *, p ≤ 0.05 (one-way analysis of variance [ANOVA], followed by Dunnett’s multiple comparisons test). **(E-G)** Effects of the truncated forms of mNRK on cell proliferation and AKT signalling. **E**) Schematic structures of the truncates. **F**) We generated Flp-In TREx 293 cell lines that expressed the truncates in a dox-dependent manner. The cells were treated with dox (1μg/mL) for two days and then subjected to the MTT assay. **G**) Cells were subjected to immunoblotting, followed by densitometric analyses. The graphs show the mean ± standard deviation (SD) of three independent experiments. ****, p≤0.0001 (unpaired two-tailed Student’s t-test).

We investigated molecular mechanisms underlying the inhibition of cell proliferation by mNRK. To identify the region(s) required for mediating the anti-proliferation effect of mNRK, we developed cells that expressed deletion mutants of mNRK in a dox-dependent manner (**Fig. 3E**). The expression of full-length mNRK and mutant #3 decreased cell numbers to a similar extent, whereas mutants #1 and #2 exerted a minor or no effect (**Fig. 3F**). Full-length mNRK and mutant #3 suppressed AKT phosphorylation, whereas mutants #1 and #2 did not (**Fig. 3G**). These results suggest that the middle region and CNH domain are required for the anti-proliferation effect of mNRK.

### NRK CNH domain acquired lipid-binding ability during mammalian evolution, thereby recruiting NRK to the plasma membrane

We then attempted to elucidate the function of the mNRK CNH domain. As mNRK localises at the plasma membrane independent of the kinase activity (Nakano et al. 2003), we speculated that the CNH domain may be important for mNRK localisation. The CNH domain of mNRK showed ∼40% amino acid similarity to the sequences of the other GCK IV family members and cNRK (**Fig. 4A**). We examined the cellular localisation of these full-length proteins and CNH domains. Although mNRK is consistently localised at the plasma membrane, mNIK, mTNIK, mMINK1, and cNRK were primarily present in the cytosol (**Fig. 4B**, upper panels). The CNH domain of mNRK was preferentially localised at the plasma membrane, but the CNH domain of NIK, TNIK, MINK1, and cNRK showed different localisation (**Fig. 4B**, lower panels). These results suggest a distinct function of the mNRK CNH domain.

**Fig. 4.**
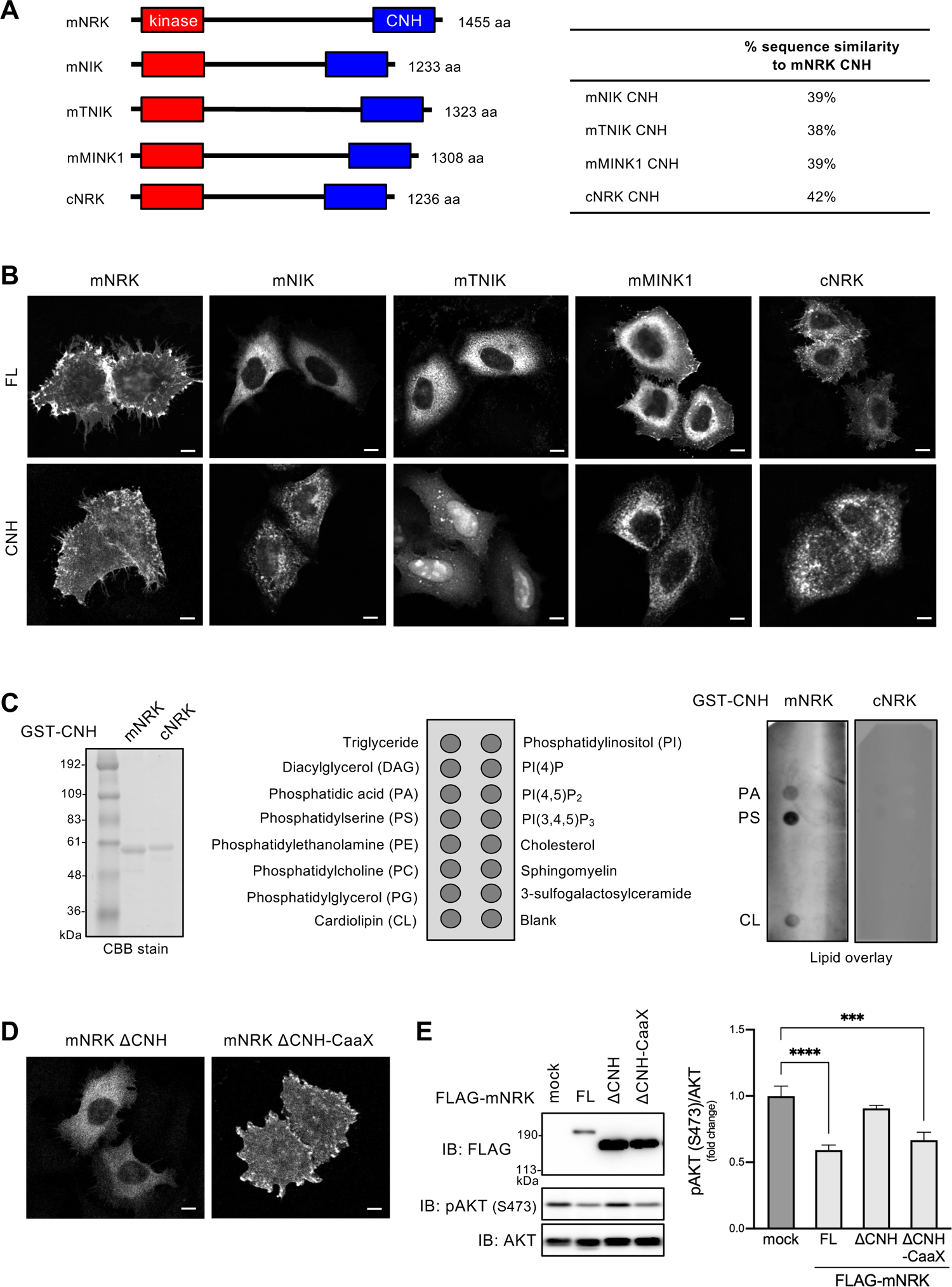
Function of the CNH domain of mNRK and related proteins. **(A)** Domain architectures of mNRK and related proteins, and sequence similarities of their CNH domains **(B)** Intracellular localisation of full length (FL) proteins of mNRK, mNIK, mTNIK, mMINK1, and cNRK and their CNH domains. HeLa cells expressing indicated proteins were subjected to immunostaining with anti-FLAG antibody. Scale bar, 10 µm. **(C)** Profiles of phospholipids that were bound to the CNH domain of mNRK. **Left panel**, CBB staining of the CNH domain of mNRK and cNRK. **Middle panel**, phospholipids spotted on the lipid strips. **Right panel**, lipid overlay assay of the CNH domain of mNRK and cNRK. The data shown are representative of two independent experiments. **(D, E)** Compensation of the deletion of the CNH domain via fusion of CaaX motif to mNRK. **D**) HeLa cells expressing indicated proteins were subjected to immunostaining. Scale bar, 10 µm. **E**, HEK293 cells expressing indicated proteins were subjected to immunoblotting, followed by densitometric analyses. The graph shows the mean ± standard deviation (SD) of three independent experiments. ***, p≤0.001; ****, p≤0.0001 (one-way analysis of variance [ANOVA], followed by Dunnett’s multiple comparisons test).

Lipid-protein overlay assay showed that the mNRK CNH domain bound strongly to phosphatidylserine (PS) and moderately to phosphatidic acid (PA) and cardiolipin (**Fig. 4C**). The cNRK CNH domain showed no detectable binding to any phospholipid (**Fig. 4C**). These results demonstrate that the mNRK CNH domain is a phospholipid-binding domain, and the NRK CNH domain acquired lipid-binding ability during mammalian evolution.

We further examined whether the CNH domain is necessary for mNRK plasma membrane localisation and whether mNRK plasma membrane localisation is important for inhibition of AKT signalling. The mNRK mutant lacking the CNH domain (ΔCNH) was present in the cytosol and did not suppress AKT phosphorylation (**Fig. 4D and E**, ΔCNH). The fusion of K-Ras prenylation motif (CaaX motif) to ΔCNH recovered the plasma membrane localisation and decreased AKT phosphorylation (**Fig. 4D and E**, ΔCNH-CaaX), indicating that the deficiency of the CNH domain was compensated by the fusion of a short plasma membrane localisation motif. These results indicate that the CNH domain recruited mNRK to the plasma membrane, thereby enabling mNRK to suppress AKT signalling.

### NRK acquired CK2-binding ability during mammalian evolution, and the region of mNRK comprising amino acids 565-868 was required for binding to CK2**β** and suppression of AKT signalling

We previously identified mNRK-interacting proteins including a CK2 catalytic subunit (CKα’) and a CK2 regulatory subunit (CK2β; Naito et al. 2020). To further elucidate the mechanisms underlying the anti-proliferating effect of mNRK, we focused on CK2 because it can modulate AKT signalling (Litchfield 2003). We performed co-immunoprecipitation and immunoblotting analysis using cells overexpressing mNRK, CK2α’, and/or CK2β. CK2α’ was co-immunoprecipitated with mNRK only in the presence of exogenous CK2β. CK2β was co-immunoprecipitated with mNRK in the presence and absence of exogenous CK2α’ (**Fig. 5A**). These results suggest that mNRK interacted with the CK2 complex by directly binding to CK2β. Next, we attempted to determine the mNRK-binding site in CK2β. The KSSR motif is one of the protein interaction motifs known in CK2β (Cao et al. 2014). The mutation in the motif (substitution of AAAA for KSSR) drastically reduced binding to mNRK (**Fig. 5B**), indicating that it is responsible for the binding.

**Fig. 5.**
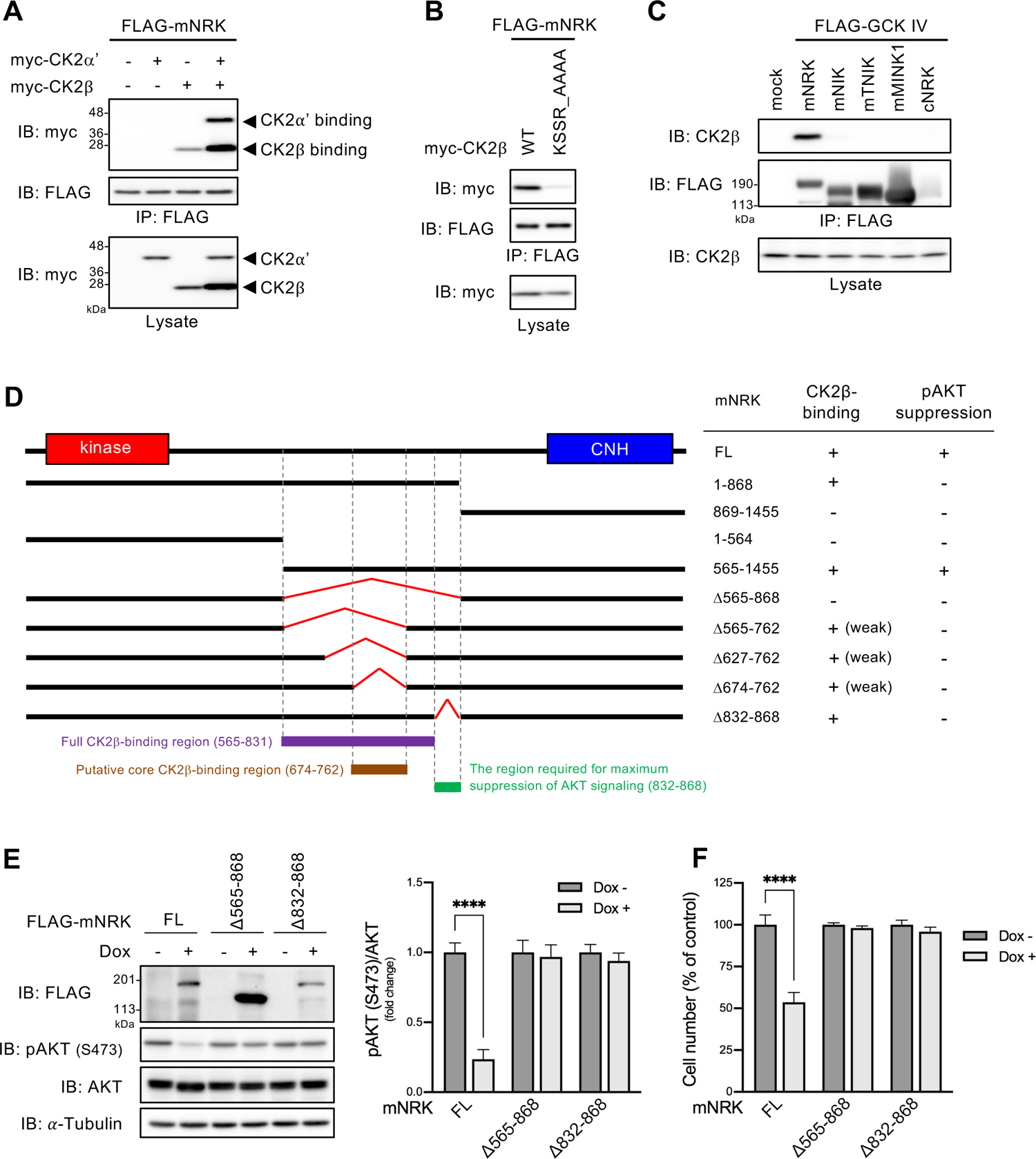
Ability of mNRK and related proteins to bind to CK2, and the relationship between mNRK binding to CK2 and its effect on AKT signalling. **(A)** Binding of mNRK to CK2β. HEK293T cells expressing indicated proteins were subjected to immunoprecipitation and immunoblotting. **(B)** mNRK binding site in CK2β. The experimental method was similar to that used for A. **(C)** No binding to CK2 was detected with mNIK, mTNIK, mMINK1, and cNRK. The experimental method was similar to that used for A. Binding of mNRK truncated mutant to CK2 and its effects on AKT signalling. **Left panel**, Schematic structure of mNRK truncated mutants. We mapped the full CK2β-binding region (aa 565-831, red), the putative core CK2β-binding region (aa 674-762, brown), and the region not necessary for binding to CK2β but required for the suppression of AKT signalling (aa 832-868, green). **Right panel**, summary of the binding of mNRK mutants to CK2β and their effects on AKT phosphorylation (See Supplementary Fig. 5A and 5B). **(E, F)** Effects of the deletion of the regions comprising aa 565-868 and aa 832-868 of mNRK on AKT signalling and cell proliferation. We generated Flp-In TREx 293 cell lines that expressed mNRK^Δ^ and mNRK^Δ^ in a dox-dependent manner. The cells were treated with dox (1μg/mL) for 2 days. **E**) The cells were subjected to immunoblotting, followed by densitometric analyses. **F**) the cells were subjected to MTT assay. The graphs show the mean ± standard deviation (SD) of three independent experiments. ****, p≤0.0001 (unpaired two-tailed Student’s t-test).

We then examined the binding ability of the other GCK IV family members and cNRK to CK2β. These proteins did not bind to CK2β (**Fig. 5C**), suggesting that NRK acquired CK2-binding ability during mammalian evolution.

To determine the CK2β-binding site in mNRK, we constructed several mNRK deletion mutants and performed co-immunoprecipitation analyses. The summary of the results is shown in **Fig. 5D**. The regions comprising amino acids (aa) 1–868 of mNRK (mNRK^1-868^) and aa 565–1455 (mNRK^565-1455^) were bound to CK2β, but the regions of aa in mNRK^869-1455^, mNRK^1-564^, and the mutant lacking aa 565–868 (mNRK^Δ565-868^) did not demonstrate any binding (**Supplementary Fig. 5A**, left panel). Hence, aa 565–868 are required for the binding activity. As described later, aa 565–762 and aa 832–868 were well conserved among eutherians. Thus, we further constructed mNRK^Δ^, mNRK^Δ^, mNRK^Δ^, and mNRK^Δ^ mutants. The mNRK^Δ^, mNRK^Δ^, and mNRK^Δ^ bound weakly to CK2β (**Supplementary Fig. 5A**, centre panel), suggesting that the region comprising aa 674– 762 was a core binding region; however, the other region weakly contributed to the binding. The mNRK^Δ832-868^ was bound to CK2β to an extent similar to that observed with full-length mNRK (**Supplementary Fig. 5A**, right panel), suggesting that the region comprising aa 832– 868 is not required for the binding.

Next, we investigated the effects of these mNRK deletion mutants on AKT phosphorylation; a summary is shown in **Fig. 5D**. The expression of mNRK^565-1455^ decreased AKT phosphorylation; however, mNRK^1-868^, mNRK^869-1455^, mNRK^1-564^ and mNRK^Δ^ expressions did not reduce the phosphorylation (**Supplementary Fig. 5B**, left panel). It should be noted that the expression level of mNRK^869-1455^ was much lower than others under this experimental condition. These results confirm that the CNH domain and the middle region comprising aa 565–868 were required for AKT suppression. The mNRK^Δ^, mNRK^Δ^, and mNRK^Δ^ did not decrease AKT phosphorylation (**Supplementary Fig. 5B**, right panel). We observed that aa 674–762, a region that is crucial for binding to CK2β, was required for AKT suppression. Surprisingly, mNRK^Δ^ did not decrease AKT phosphorylation (**Supplementary Fig. 5B**, right panel), suggesting that the region comprising aa 832-868 was required for AKT suppression despite not contributing toward binding to CK2β.

We then generated cells with dox-induced expression of mNRK^Δ565-868^ and mNRK^Δ868^. As expected, the expression of these mutants did not decrease AKT phosphorylation (**Fig. 5E**) and cell proliferation (**Fig. 5F**).

### Region comprising amino acids 565–868 of mNRK inhibited CK2 activity *in vitro*

Next, we investigated whether mNRK^565-868^ inhibited CK2 activity *in vitro* (**Fig. 6A**). A CK2 substrate protein PTEN^190-403^ (mutant lacking N-terminal phosphatase domain) was phosphorylated by a commercial CK2 complex in the presence or absence of mNRK^565-868^ (**Fig. 6A**). The phosphorylation levels were measured by immunoblotting with phospho-PTEN (S380) antibody and phospho-CK2 substrate antibody. The result showed that mNRK^565-868^ inhibited CK2 activity (**Fig. 6B**). We named this region the CK2-inhibitory region (CIR).

**Fig. 6.**
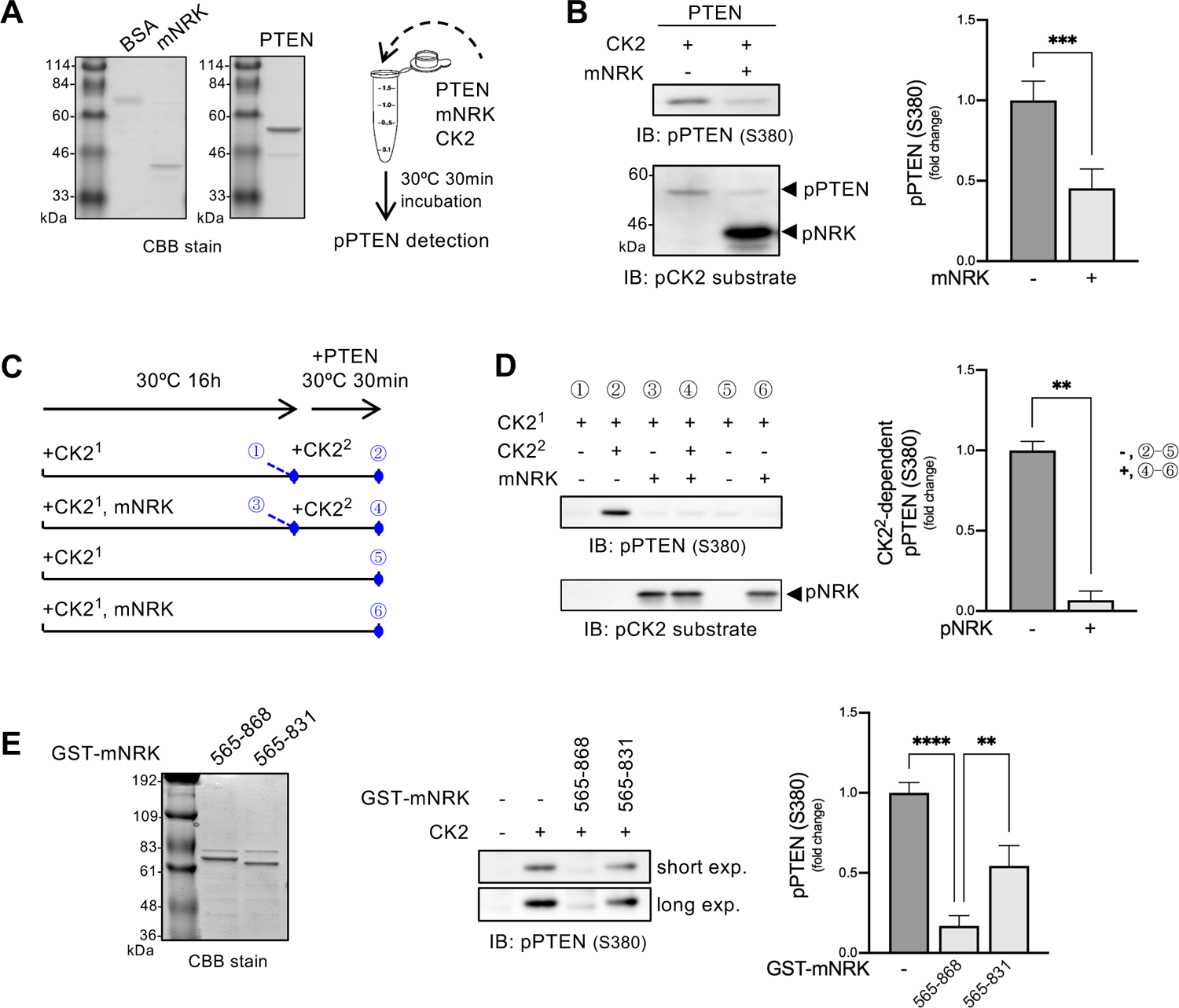
Effects of the middle region of mNRK on CK2 kinase activity. **(A)** CBB staining of purified mNRK^565-868^ and GST-PTEN^190-403^ and schematic representation of *in vitro* kinase assay in **B**. **(B)** Inhibition of CK2 activity by mNRK^565-868^. CK2, GST-PTEN^190-403^, and mNRK^565-868^ were mixed in a kinase assay buffer containing ATP and incubated at 30 °C for 30 min. The samples were then subjected to immunoblotting, followed by densitometric analyses. The graph shows the mean ± standard deviation (SD) of four independent experiments. ***, p≤0.001 (unpaired two-tailed Student’s t-test). **(C)** Schematic representation of the *in vitro* kinase assay and the sampling time of each sample (#1–6) shown in **D**. **(D)** mNRK^565-868^ inhibition of CK2 activity via a non-competitive mechanism. mNRK^565-868^ was phosphorylated by preincubation with CK2 (denoted as CK2^1^) in a kinase assay buffer containing ATP at 30 °C for 16 h. The samples were then mixed with CK2 (denoted as CK2^2^), GST-PTEN^190-403^, and ATP and incubated at 30 °C for 30 min, followed by immunoblotting and densitometric analyses. We subtracted the value of sample #5 from sample #2 and sample #6 from sample #5, marking them as those with phosphorylation of PTEN by CK2^2^ in the absence or presence of phospho-NRK in the graph, respectively. The graph shows the mean ± standard deviation (SD) of four independent experiments. **, p≤0.01 (unpaired two-tailed Student’s t-test). **(E)** Importance of the region comprising aa 832-868 for maximum inhibition of CK2 activity by mNRK^565-831^. CK2, GST-PTEN^190-403^, and GST-mNRK (mNRK^565-868^ or mNRK^565-831^) were mixed in a kinase assay buffer containing ATP and incubated at 30 °C for 30 min. The samples were then subjected to immunoblotting, followed by densitometric analyses. The graph shows the mean ± standard deviation (SD) of three independent experiments. **, p≤0.01; ****, p≤0.0001 (one-way analysis of variance [ANOVA], followed by Tukey’s multiple comparisons test).

We also observed CK2-dependent phosphorylation of mNRK^565-868^ (**Fig. 6B**). This raised the possibility that the CIR may compete with PTEN as a substrate for CK2. To test this hypothesis, we prepared phospho-mNRK^565-868^ by preincubation with CK2 (denoted as CK2^1^) and examined its effect on the activity of CK2 that was added later (denoted as CK2^2^; **Fig. 6C**). The results show that PTEN^190-403^ was strongly phosphorylated by CK2^2^ in the absence of phospho-mNRK^565-868^ (**Fig. 6D**, #1 vs #2). However, it was phosphorylated to a small extent by CK2^2^ in the presence of phospho-mNRK^565-868^ (#3 vs #4). CK2^1^ added at the start of the 16 h preincubation showed little activity in the subsequent 30-min incubation (#5 and #6), probably because CK2^1^ might have lost its activity during long-term preincubation. We also found similar phosphorylation levels of mNRK^565-868^ in samples #3, #4, and #6 (**Fig. 6D**, bottom panel), confirming that almost all mNRK^565-868^ was phosphorylated by CK2^1^.

Densitometric analyses showed that phospho-mNRK^565-868^ inhibited CK2^2^-dependent phosphorylation of PTEN^190-403^ (**Fig. 6D**, right panel). These data suggest that the CIR inhibited CK2 activity via a non-competitive mechanism.

In **Fig. 5**, we show that the region comprising aa 832–868 of mNRK did not contribute toward binding to CK2β but was essential for AKT suppression. We examined the effects of GST-tagged mNRK^565-868^ and mNRK^565-831^ (lacking the region comprising aa 832–868) on CK2 activity. The results show that mNRK^565-868^ strongly decreased CK2-dependent phosphorylation of PTEN^190-403^, whereas mNRK^565-831^ only demonstrated a minor reduction in phosphorylation (**Fig. 6E**). These results indicate that the region of mNRK comprising aa 832–868 plays an important role in the inhibition of CK2 activity.

### mNRK suppressed AKT signalling by modulating the CK2-PTEN-AKT pathway

We then examined whether mNRK inhibited CK2 activity in cells. Dox-induced mNRK expression reduced PTEN S380 phosphorylation (**Fig. 7A**). The mNRK expression did not affect the protein levels of each CK2 subunit and the interaction of CK2α’ with CK2β (**Fig. 7A**). These results indicate that mNRK inhibited cellular CK2 activity without affecting the formation of the CK2 complex.

**Fig. 7.**
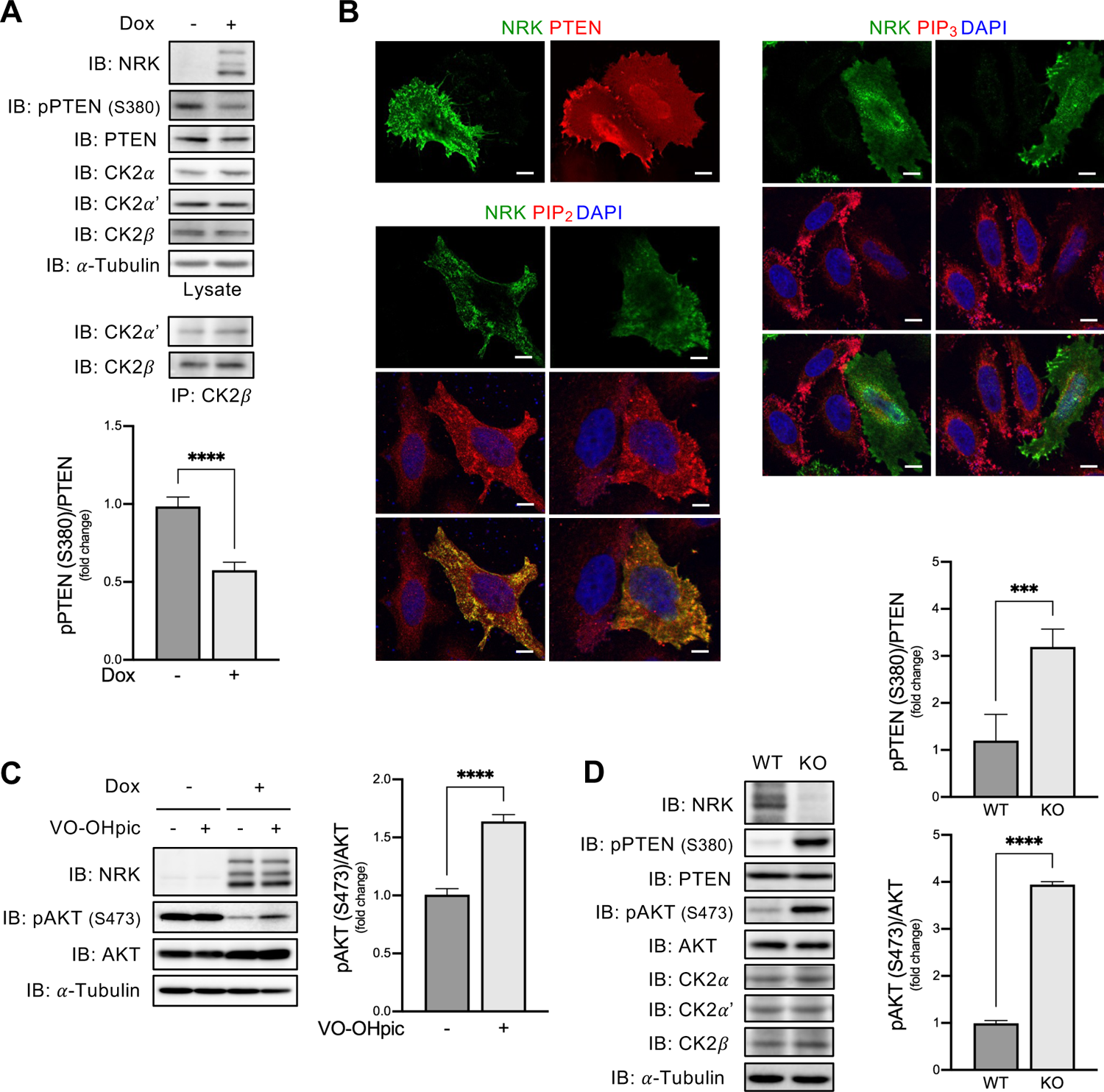
Effects of mNRK on the CK2-PTEN-AKT pathway. **(A)** Effects of mNRK on PTEN phosphorylation in cultured cells. Flp-In TREx 293 cells with dox-inducible expression of mNRK were treated with dox (1 μg/mL) for 2 days. The cells were subjected to immunoblotting, followed by densitometric analyses. The graph shows the mean ± standard deviation (SD) of four independent experiments. ****, p ≤ 0.0001 (unpaired two-tailed Student’s t-test). **(B)** Effects of mNRK on PTEN localisation and levels of PIP_2_ and PIP_3_. **Upper left panel**, HeLa cells expressing myc-mNRK and FLAG-PTEN were subjected to cell staining with anti-myc antibody (green) and anti-FLAG antibody (red). **Lower left and right panels**, cells expressing myc-mNRK were subjected to staining with anti-myc antibody (green), anti-PIP_2_ or PIP_3_ antibodies (red), and DAPI (blue). Scale bars, 10 μ **(C)** Restoration of AKT phosphorylation by PTEN inhibitor in mNRK-expressing cells. Flp-In TREx 293 cells with dox-inducible expression of mNRK were treated with dox (1μg/mL) for 2 days and treated with a PTEN inhibitor VO-OHpic (1 μ The cells were subjected to immunoblotting, followed by densitometric analyses. The graph shows the mean ± standard deviation (SD) of four independent experiments. ****, p≤0.0001 (unpaired two-tailed Student’s t-test). **(D)** Effects of mNRK on PTEN phosphorylation in mouse placentas. Placenta tissues were lysed at 18.5 dpc and subjected to immunoblotting with indicated antibodies, then densitometric analyses. The graph shows the mean ± standard deviation (SD; WT, n = 5; KO, n = 5). ***, p≤0.001, ****, p≤0.0001 (unpaired two-tailed Student’s t-test).

We examined the effect of mNRK expression on PTEN localisation. PTEN was detected in the cytosol and nucleus in mNRK-negative cells. A part of PTEN was also detected at the plasma membrane in mNRK-positive cells (**Fig. 7B**). The mNRK expression increased PIP_2_ and decreased PIP_3_. These observations suggest that mNRK inhibited CK2-mediated phosphorylation of PTEN, thereby mediating the localisation of PTEN on the plasma membrane and PIP_3_ dephosphorylation.

Next, we examined whether mNRK-induced AKT suppression was mediated by PTEN. We used cells with or without dox-inducible expression of mNRK and treated them with PTEN inhibitor VO-OHpic. The PTEN inhibitor showed minor effects on AKT phosphorylation in cells without mNRK expression. Dox-induced mNRK expression decreased AKT phosphorylation; PTEN inhibitor restored it (**Fig. 7C**). These results suggest that PTEN is mostly inactive in cells without mNRK expression and that NRK activates PTEN, thereby leading to AKT suppression.

Next, we examined the role of mNRK in the regulation of the CK2-PTEN-AKT pathway under *in vivo* conditions using NRK knockout (KO) mice. Consistent with earlier reports, KO placentas did not express mNRK and were larger than WT placentas. KO placentas showed increased phosphorylation levels of PTEN S380 as well as AKT S473 compared with WT placentas (**Fig. 7D**). These results suggest that mNRK inhibits phosphorylation of PTEN by CK2 and activates PTEN, leading to the suppression of AKT signalling in the placenta.

### Functional regions in eutherian NRK underwent negative selection

Given that mNRK plays an important role in regulating placental development, we speculated that its functional regions would be affected by negative selection in eutherians. To evaluate selective pressure during *Nrk* evolution, we compared coding sequences of fifteen eutherian *Nrk* orthologues and calculated the ratio of non-synonymous to synonymous substitution rates (dN/dS). The kinase and CNH domains showed low dN/dS values (0.14 and 0.20 on average, respectively; dN/dS < 1; **Fig. 8A**), suggesting that they are under strong purifying selection. The amino acid sequence alignment of the CNH domain in vertebrate NRK orthologues shows a similarity in eutherian sequences and dissimilarity among eutherian and other sequences (**Fig. 8B**), consistent with the dN/dS and evolutionary rate analysis (**Fig. 2**).

**Fig. 8.**
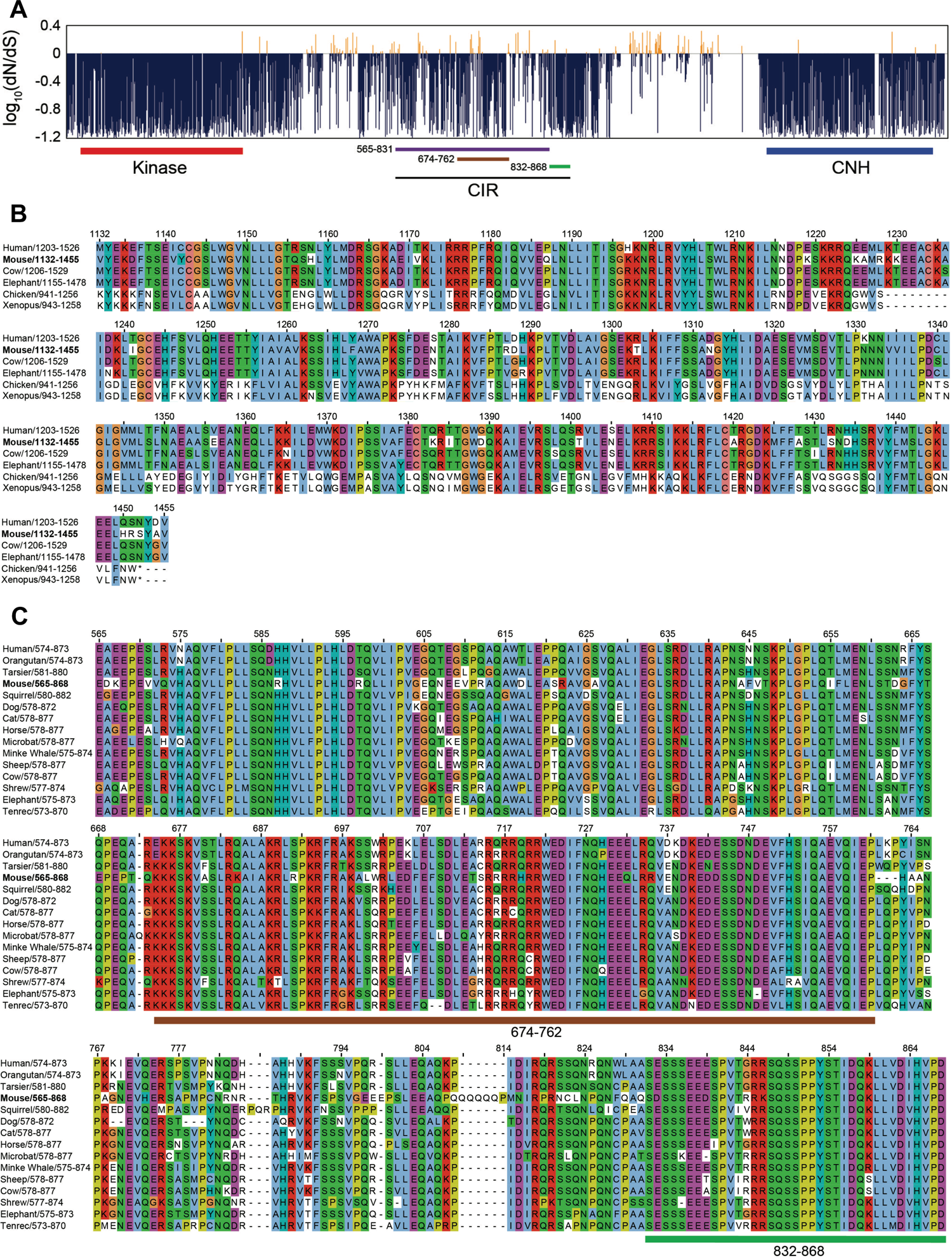
Evolutionary pressure on mNRK functional regions. **(A)** Negative selection of eutherian *Nrk* sequences. The graph shows the ratio of non-synonymous per synonymous substitution (dN/dS) estimated using 15 eutherian *Nrk* sequences; dN/dS<1 predicts negative selection. Functional domains and regions are indicated: 565-831, full CK2β binding region; 674-762, putative core CK2β-binding region; 832-868, the region required for the maximum inhibition of CK2 activity. **(B)** Conservation of amino acid sequences of NRK CNH domains among eutherians. We selected eutherians (human, mouse, cow, elephant) and non-mammals (chicken and *Xenopus*), aligned their NRK CNH domain sequences, and coloured them according to the ClustalX colour scheme. **(C)** Conservation of amino acid sequences of NRK CK2-binding regions among eutherians. We selected 15 eutherians, aligned their regions orthologous to the mNRK CK2-inhibitory region, and coloured them according to the ClustalX colour scheme.

The CIR showed low dN/dS values (**Fig. 8A**). We performed amino acid sequence alignment of the CIR. The sequences of non-eutherian NRK could not be aligned owing to their much lower similarity to mNRK. Two functional regions were examined; a region (aa 674–762; dN/dS = 0.38) important for binding to CK2β, and a region (aa 832–868; dN/dS = 0.23) not required for binding to CK2β but necessary for maximum inhibition of the CK2 activity. As expected, these two regions appear to be well conserved among eutherians (**Fig. 8C**).

## Discussion

This study showed that the eutherian *Nrk* gene underwent multi-scale evolution in terms of genomic location, exon structure, sequence of the protein-coding region, and gene expression regulation. NRK underwent extensive amino acid substitutions in the ancestor of placental mammals and has been conserved since then. Through this, eutherian NRK seems to have acquired the lipid-binding CNH domain and the CIR. We also found that mNRK inhibits placental cell proliferation by modulating the CK2-PTEN-AKT pathway, and this function can be attributed to this molecular evolution (**Fig. 9**, working hypothesis).

**Fig. 9.**
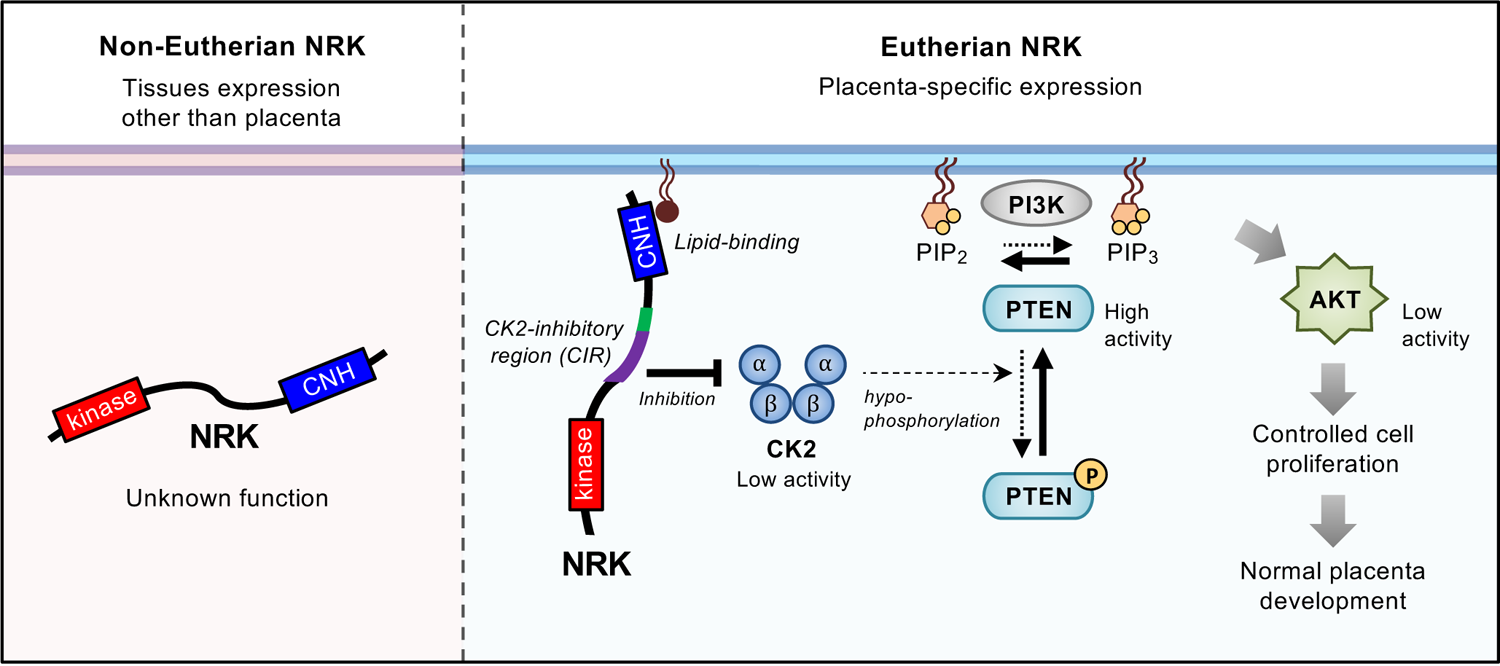
Model of molecular mechanisms underlying inhibition of placental cell proliferation by eutherian NRK and its molecular evolution to acquire this function. NRK is highly expressed in the placenta and underwent rapid molecular evolution during early mammalian evolution. The changes in amino acid sequences in the CNH domain and the middle region conferred plasma membrane localisation and CK2 inhibition activities, respectively. Eutherian NRK is localised at the plasma membrane and inhibited CK2, thereby mitigating CK2-dependent inhibition of PTEN, leading to the suppression of AKT signalling and cell proliferation. Through these mechanisms, eutherian NRK played an essential role in normal placental development by preventing hyperplasia of the placenta.

### Evolutionary history of *Nrk* leading to an X-linked placenta-specific gene

Mouse and human *Nrk* genes are located on Xq and are descended from proto-X chromosome orthologous to chicken 4p. A previous report indicated that approximately 30 subregions with sequence homology were observed upon comparing human Xq with chicken 4p. However, their positions and directions differed, implying many recombination events during the evolution of their common ancestor (Ross et al. 2005). Our result suggests that the genomic region containing *Nrk* was relocated owing to multiple genomic recombinations during the early evolution of mammals.

Consistent with a theory that selection favours an increase in the frequency of genes with sexually antagonistic traits on the X chromosome (Rice 1984), the mouse X chromosome is enriched for genes preferentially expressed in the placenta (Khil et al. 2004). Our study added eutherian *Nrk* to the list of X-linked placenta-specific genes. Recently, Liu et al. screened the eutherian X chromosome to detect the regions with accelerated evolution in the ancestral lineage of eutherians and found that the region with the highest accelerated evolution is located in the coding regions of *Nrk* (Liu et al. 2021). Our study reveals that *Nrk* underwent accelerated evolution in terms of exon gain/loss, sequence insertion events in the eutherian and therian ancestors, and rapid sequence substitutions in the eutherian ancestor. Accelerated evolution has been frequently observed in duplicated genes such as those in teleost fish (Brunet et al. 2006). However, we found no paralogous gene of *Nrk* duplicated in or before the eutherian ancestor, suggesting that the eutherian-specific rapid evolution of *Nrk* was not caused by gene duplication. Although the expression of *Nrk* genes in non-eutherians and the function of their coding proteins are largely unclear, *Nrk* seems to have been co-opted to generate eutherian placenta. We speculate that *Nrk* became a placental-specific gene in the eutherian or therian ancestor and was subsequently subjected to strong positive evolutionary pressure to optimise its protein function in the placenta. After optimisation, it was subjected to negative evolutionary pressure to maintain the function (**Fig. 8A**).

Many proteins are specifically expressed in the placenta and contribute to placental function and regulation (Rossant and Cross 2001; Rawn and Cross 2008; Woods et al. 2018). They can be divided into two groups, namely, those found only in a few mammalian species and those widely shared among mammals. NRK is included in the latter group. Most proteins in the former group may enable diversity among species in the placental morphology and mother-foetus interactions. In contrast, proteins in the latter group are thought to be involved in the fundamental development, function, and regulation of the placenta. The evolutionary history of the latter group can be further classified into several patterns. The first pattern involves proteins encoded by genes that were newly acquired during early mammalian evolution. For example, a placenta-specific secreted protein PLAC1 is not found in marsupials or monotremes but is found in eutherians (Devor 2014). A retrotransposon-derived PEG10 is found only in placental mammals (Suzuki et al. 2007). The next pattern involves those present in many taxons of vertebrates, while they have come to be expressed in the placenta during early mammalian evolution, without changing their primary protein function. For example, HAND1, a transcription factor found in vertebrates, regulates gene expression in neurons in chickens (Howard et al. 1999). It is also expressed in the placenta in mice and thought to regulate placental gene expression as transcriptional factor (Riley et al. 1998). NRK does not fit into these patterns; it was present in mammalian ancestors but acquired new functional domains/regions during early mammalian evolution, changing its molecular functions to regulate placental development. To our knowledge, no placenta-specific protein other than NRK is known with such an evolutionary history.

### Unique role of eutherian NRK in regulating AKT signalling among GCK IV kinases

We also found that other members of GCK IV kinases (mNIK, mTNIK, mMINK1) and cNRK were not localised at the plasma membrane and did not interact with CK2 or inhibited AKT signalling under our experimental conditions. NIK is reported to enhance the invasion of some cancer cells (Wright et al. 2003). TNIK potentiates WNT and AKT signalling to promote tumorigenesis (Yu et al. 2014; Masuda et al. 2016). MINK1 interacts with mTORC2 to enhance AKT signalling and promote cancer cell metastasis (Daulat et al. 2016). While the function of non-eutherian NRK remains unclear, these facts demonstrate that, among all GCK IV kinases, only eutherian NRK functions as a negative regulator of AKT signalling.

### Roles of the kinase domain, CNH domain, and CIR of mNRK

This study shows distinct functions of the kinase domain, CNH domain, and CIR mNRK. The kinase domain of eutherian NRK underwent negative selection (**Fig. 8A**), indicating a crucial function mediated by this domain. mNRK phosphorylates cofilin, thereby enhancing actin polymerisation (Nakano et al. 2003). mNRK can also activate the JNK pathway in a kinase activity-dependent manner (Nakano et al. 2000; Kakinuma et al. 2005). However, in this study, the kinase domain was not required for mNRK-induced inhibition of AKT signalling or cell proliferation. We speculate that NRK may inhibit AKT signalling, possibly not via mechanisms involving actin polymerisation or JNK activation.

The CNH domain contributed to recruiting mNRK to a site near the plasma membrane. The localisation of mNRK on the plasma membrane was required to inhibit AKT signalling. The CNH domain was bound to phospholipids, although it was unclear whether this binding activity was sufficient for its localisation on the plasma membrane. Further study is required to investigate the detailed mechanisms. Researchers have reported several phospholipid-binding domains, most of which are well conserved among eukaryotes (Lemmon 2008). To our knowledge, this is the first report of the mammalian-specific intracellular phospholipid-binding domain. The CNH domain of NRK may be useful to study the molecular evolution of the phospholipid-binding domain.

The CIR, the region comprising aa 565–868 of mNRK, was bound to the CK2β regulatory subunit and inhibited the kinase activity of the CK2 complex via a non-competitive mechanism. In the CIR, the region comprising aa 565–831 is responsible for binding to CK2β. CK2β is thought to recruit substrates and/or regulators of the CK2 complex (Bibby and Litchfield 2005). The region comprising aa 565–831 weakly inhibited the CK2 activity *in vitro* (**Fig. 6E**), indicating that this region may impair the ability of CK2β to recruit substrates to the CK2α/α’ catalytic subunit. We also found that the region comprising aa 832–868 was not required for binding to CK2β; however, it was required for the maximal inhibition of the CK2 activity (**Fig. 5D** and **Fig. 6E**). The region comprising aa 832–868 has putative phosphorylation sites for CK2, but how this region is involved in the inhibition remains unclear. CK2 is known to be inhibited by several proteins (Homma et al. 2002; Llorens et al. 2003), including the intrinsically disordered protein Nopp140. Nopp140 primarily binds to CK2β and is phosphorylated by the CK2 complex (Li et al. 1997). The phosphorylated sequence in Nopp140 binds to the active site of CK2α, thereby inhibiting the kinase activity (Lee et al. 2013). These studies suggest that Nopp140 may have different regions that interact with CK2α and CK2β, and the two regions work in concert to inhibit CK2 effectively. We /α’ and /α’ interaction region. Similar to Nopp140, a large part of the CIR is predicted to be intrinsically disordered (Naito et al. 2020). In general, the disordered regions of proteins appear to evolve rapidly owing to relaxed purifying selection (Brown et al. 2011). We speculate that the disordered regions in the middle of NRK underwent rapid evolution in the ancestor of placental mammals and became the CIR.

### Molecular mechanism underlying the inhibition of AKT signalling by mNRK

We observed that mNRK overexpression in cultured cells inhibited the PTEN phosphorylation activity of CK2. Based on previous studies, mNRK must mitigate CK2-dependent negative regulation of PTEN (Vazquez et al. 2000; Torres and Pulido 2001; Rahdar et al. 2009). Consistently, mNRK overexpression decreased PIP_3_ levels accompanied by an increase in PIP_2_ levels, leading to the suppression of AKT signalling and cell proliferation. In addition to phosphorylating PTEN, CK2 can directly phosphorylate AKT S129 and thereby enhance AKT kinase activity (Risso et al. 2015). CK2 may also inhibit phosphatases targeting phospho-T308 and phospho-S473 of AKT, leading to the enhancement of AKT kinase activity (Trotman et al. 2006; Chatterjee et al. 2013). Therefore, it cannot be denied that additional mechanisms are involved in the attenuation of AKT signalling by mNRK.

### Mechanism for the regulation of placental development

We found that *Nrk* deficiency increased PTEN as well as AKT phosphorylation in mouse placenta. CK2, PTEN, and AKT have been implicated in the regulation of placental development. CK2 is expressed in the placenta from early pregnancy, and is required for cell proliferation, migration, and differentiation (Abi Nahed et al. 2020). PTEN^+/-^ mice showed an enlarged spongiotrophoblast layer and approximately 20% increase in placental weight (Church et al. 2012), similar to NRK KO mice (Denda et al. 2011), albeit with a milder phenotype. Phosphatase of regenerating liver (PRL)-2 is a protein responsible for PTEN degradation, and PRL-2 KO mice showed a small spongiotrophoblast layer (Dong et al. 2012). AKT1 deficiency reduced the weight of mouse placenta (Yang et al. 2003), and an AKT inhibitor inhibited placental cell proliferation (Morioka et al. 2017). Based on the evidence, we believe that NRK prevents hyperplasia of the placenta by modulating the CK2-PTEN-AKT pathway. So far, the types of physiological situations in which AKT signalling is regulated via the CK2-PTEN-AKT pathway remain unclear. Given that *Nrk* expression is pronounced in the placenta during late pregnancy, our study suggests that tissue-specific expression of mNRK is the first mechanism to regulate AKT signalling via the CK2-PTEN-AKT pathway.

### Limitations of this study and future directions

The data on tissue expression patterns of the *Nrk* gene in vertebrates were limited. In order to better understand the involvement of *Nrk* in placental evolution, it is important to investigate their expression in tissues that supply nutrients and oxygen to the foetus, such as the primitive placenta in marsupials, the chorioallantoic membrane in reptiles and birds, and placenta-like structures in some fish. (2) It is also important to identify the regulatory mechanism underlying placenta-specific expression of eutherian *Nrk g*ene. Multiple recombination events in the genomic region containing the *Nrk* gene occurred during early mammalian evolution. This might have led to the interchange of transcriptional regulatory elements between genes, resulting in placenta-specific expression of the *Nrk* gene. (3) We investigated the molecular mechanism by which mNRK suppresses AKT signalling using HEK293-derived cells. However, *in vivo*, mNRK is highly expressed in placental spongiotrophoblasts. As spongiotrophoblasts specifically express some proteins that regulate AKT signalling (Takao et al. 2012), the molecular mechanism may be more complicated in these cells. (4) The organisation of eutherian placentas is diverse, and it is unclear whether non-mouse placenta possesses cells similar to mouse spongiotrophoblasts. Recent single-cell RNA-seq analysis showed that human *NRK* is expressed in placental syncytiotrophoblasts and cytotrophoblasts (Vento-Tormo et al. 2018). The well-conserved amino acid sequences of the functional domains/regions of eutherian NRK imply their conserved molecular functions. It is important to investigate how NRK functions in the placenta of species other than mice.

## Conclusions

This is the first report suggesting that NRK prevents over-proliferation of placental cells by modulating the CK2-PTEN-AKT pathway. Reduced NRK levels, the loss of NRK function due to mutations in the CNH domain or the CIR, or abnormalities in the downstream pathway of NRK may lead to pregnancy complications. We also provided evidence for the unique and rapid molecular evolution of NRK, during which it acquired the anti-proliferation function. We believe that NRK evolution facilitated the proper control of placenta development in mammals.

## Acknowledgments

We would like to thank the Biomaterials Analysis Division, Tokyo Institute of Technology, for DNA sequencing analysis, confocal microscopy analysis, and mice breeding. We thank Dr. Hiroshi Iwasaki and Dr. Hitoshi Nakatogawa (Tokyo Institute of Technology) for helpful discussions. We thank Dr. Mikiko Tanaka for the chicken embryo DNA library. We thank Dr. Yuka Madoka for help with the plasmid preparation. This work was supported by the Sasakawa Scientific Research Grant from the Japan Science Society (AE, grant number: 2018-4012) and the Takeda Science Foundation (TF). We thank Editage (www.editage.com) for English language editing.

---

**Supplementary Figure 1.**
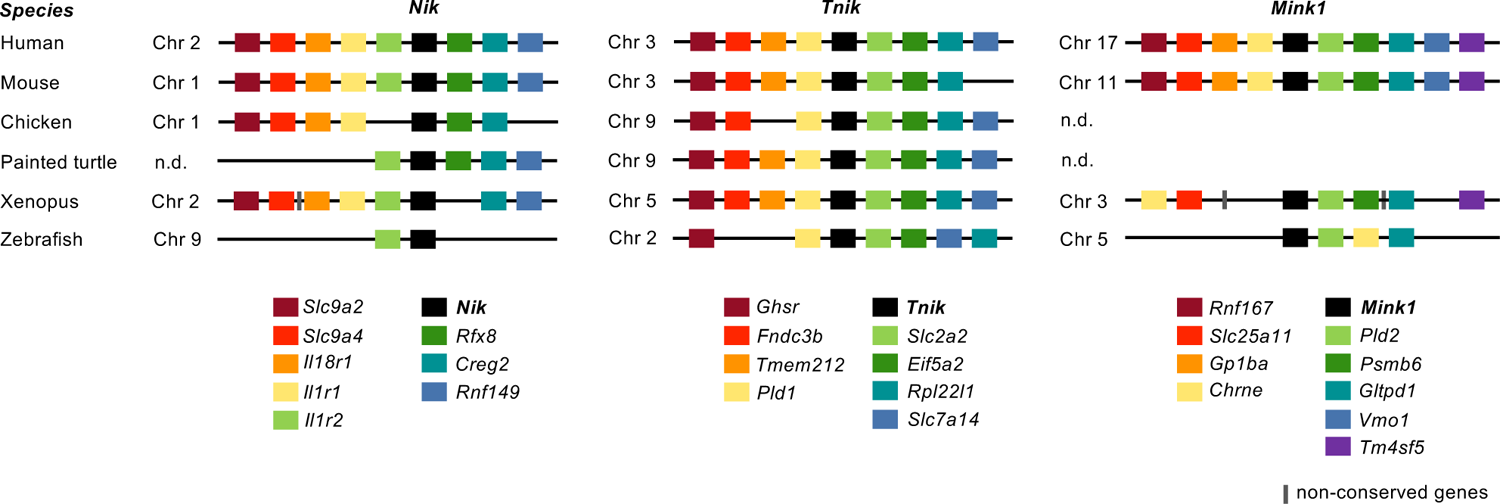
Synteny analysis of the *Nik, Tnik, and Mink1* genes Synteny conservation of the region surrounding the *Nik, Tnik, and Mink1* genes. Gray boxes indicate non-conserved genes.

**Supplementary Figure 2.**
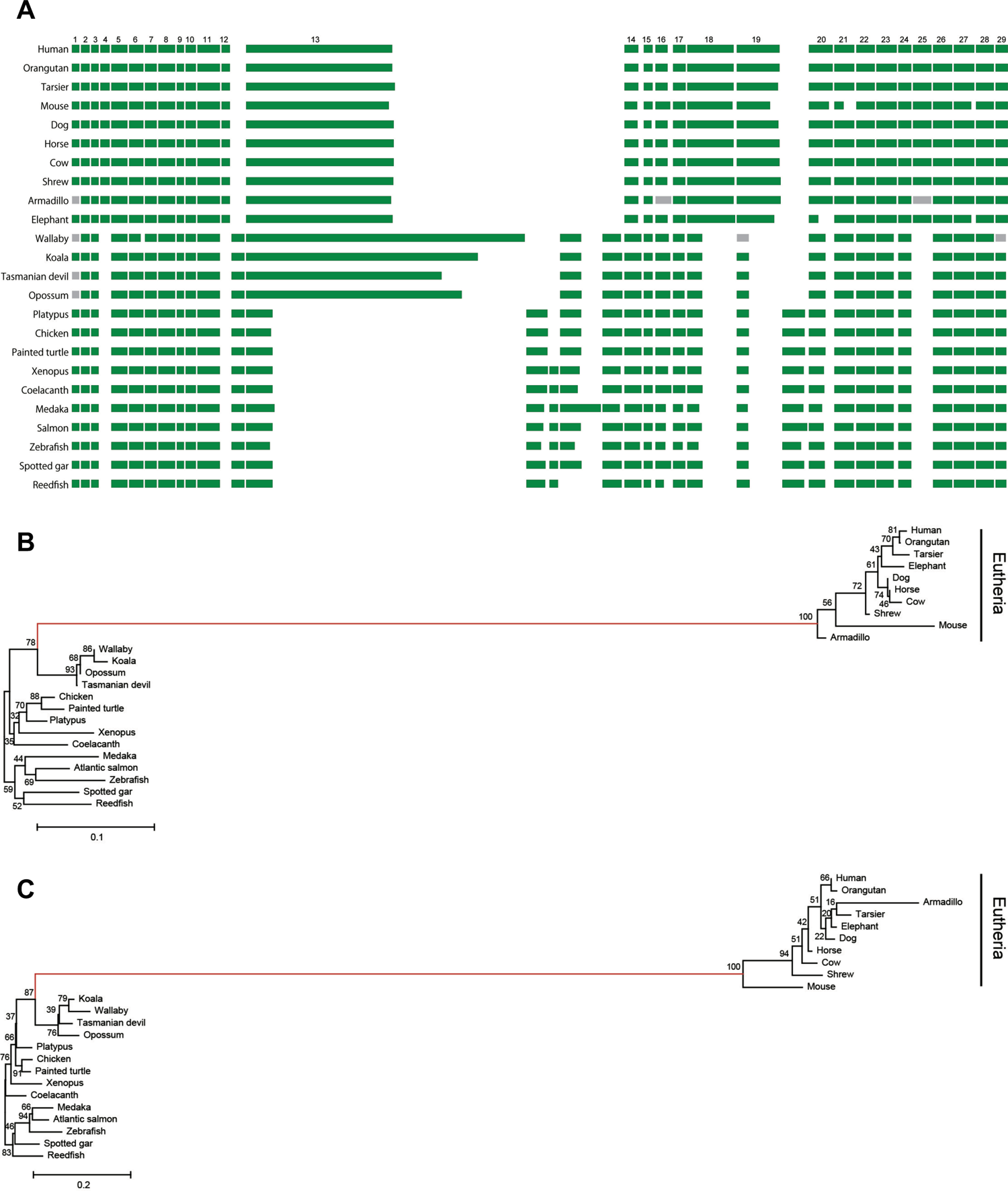
Exon structures of the *Nrk* gene and neighbour-joining tree for the amino acid sequences of the NRK domains **(Related to** Fig. 2**) (A)** Exon structures of vertebrate *Nrk*. Green boxes correspond to the coding exons, while grey boxes indicate undetermined exons. All boxes are arranged left for each exon. The numbers above represent exon numbers in eutherians. **(B)** Neighbour-joining (NJ) tree of the kinase domain corresponding to exons 1–11 of *Nrk*, excluding the eutherian-specific exon 4. Numbers on branches are node support based on 1,000 bootstrap replicates. **(C)** NJ tree of the CNH domain corresponding to exons 21–29 of *Nrk*, excluding the eutherian-specific exon 25.

**Supplementary Figure 3.**
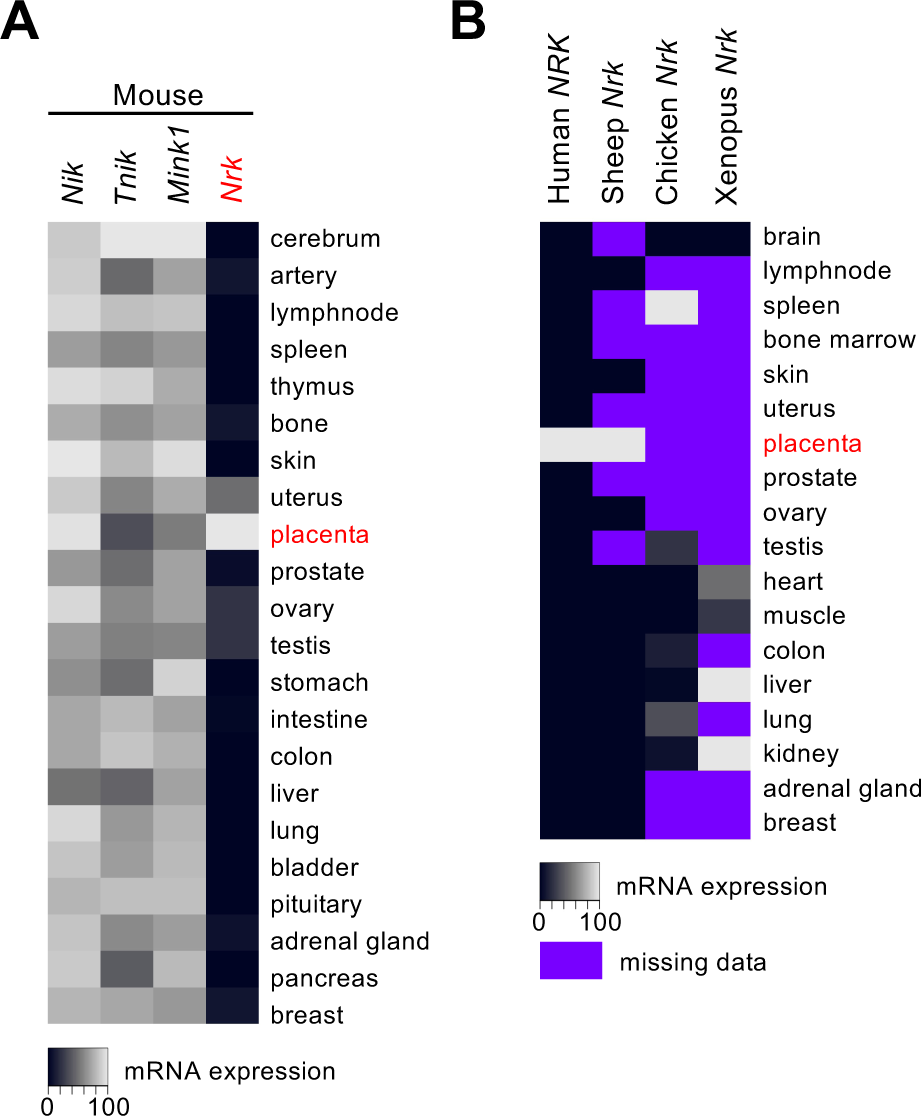
Tissue expression patterns of vertebrate *Nrk* **(A)** Heatmap of *Nrk* gene expression in different mouse tissues in comparison to the expression patterns of the other GCK IV family members (*Nik, Tnik, Mink1*). The colour intensity reflects the mRNA expression levels from the lowest (black) to the highest (white). **(B)** Heatmap of *Nrk* gene expression in human, sheep, chicken, and *Xenopus* tissues. The deep purple panels represent the missing data.

**Supplementary Figure 4.**
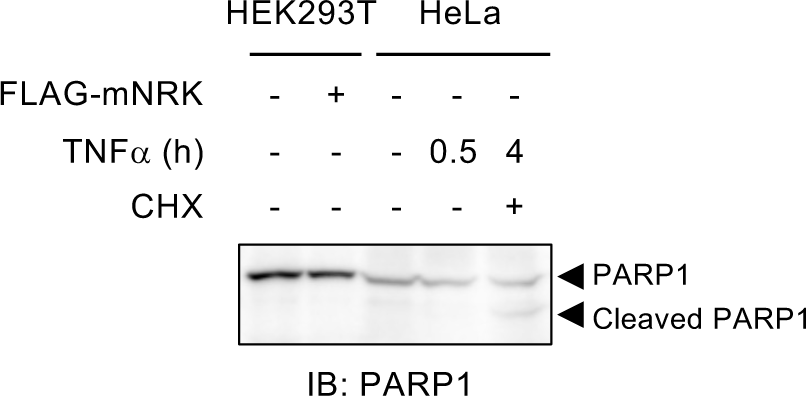
Effects of mNRK on cell apoptosis **(Related to** Fig. 3) HEK293T cells were transfected with plasmid encoding FLAG-mNRK. As a positive control, HeLa cells were stimulated with TNF (10 ng/mL for 0.5 or 4 h) and CHX (10 µg/mL for 4 h). The lysates were subjected to immunoblotting using an anti-PARP1 antibody.

**Supplementary Figure 5.**
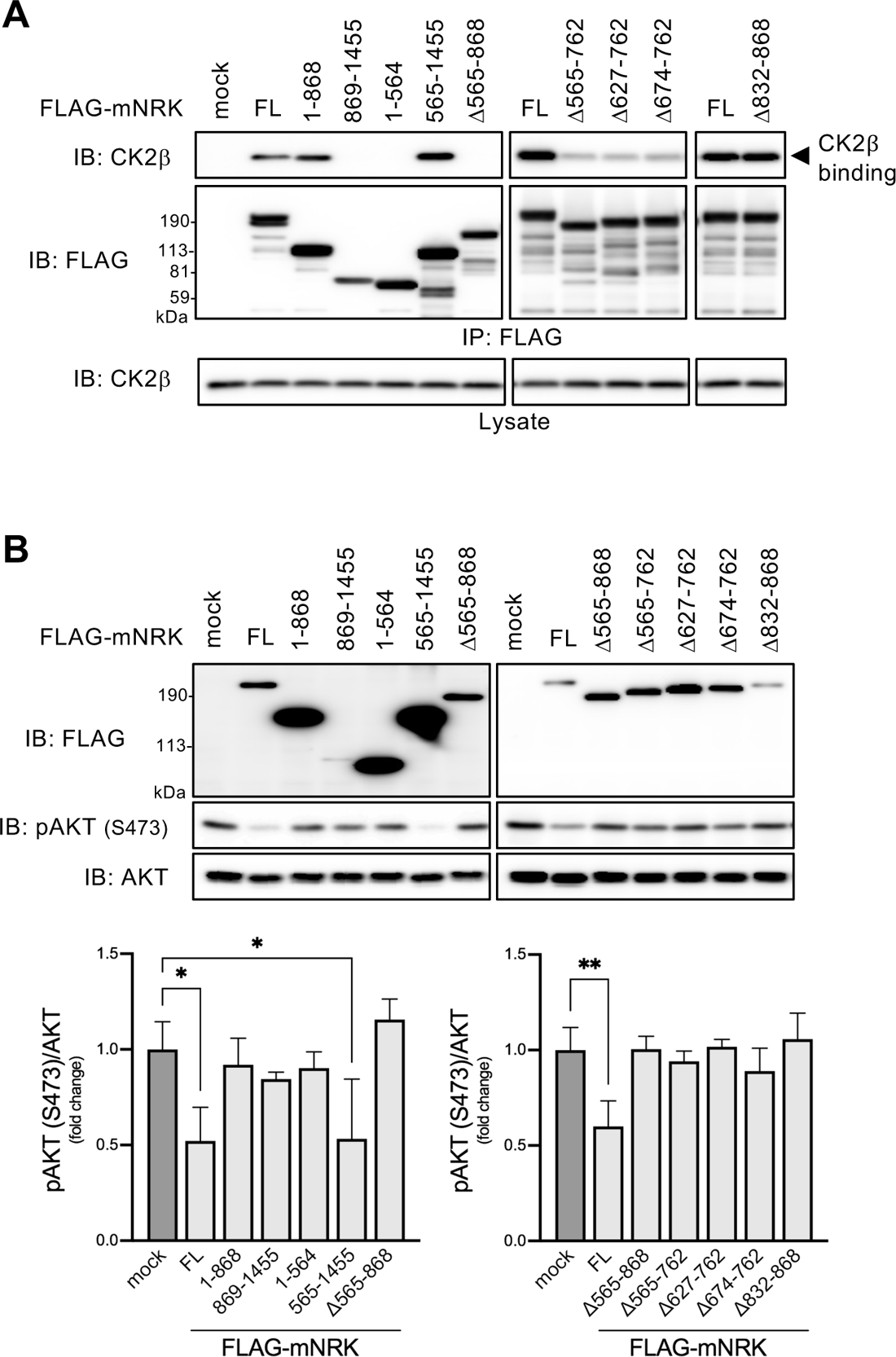
Binding of mNRK truncated mutant to CK2 and its effects on AKT signalling **(Related to** Fig. 5**) (A)** Regions in mNRK responsible for binding to CK2β. HEK293T cells expressing indicated proteins were subjected to immunoprecipitation and immunoblotting. **(B)** Regions in mNRK responsible for the suppression of AKT phosphorylation. HEK293 cells expressing indicated proteins were subjected to immunoblotting, followed by densitometric analyses. The graph shows the mean ± standard deviation (SD) of three independent experiments. *, p≤0.05; **, p≤0.01 (one-way analysis of variance [ANOVA], followed by Dunnett’s multiple comparisons test).

**Supplementary Table 1.**
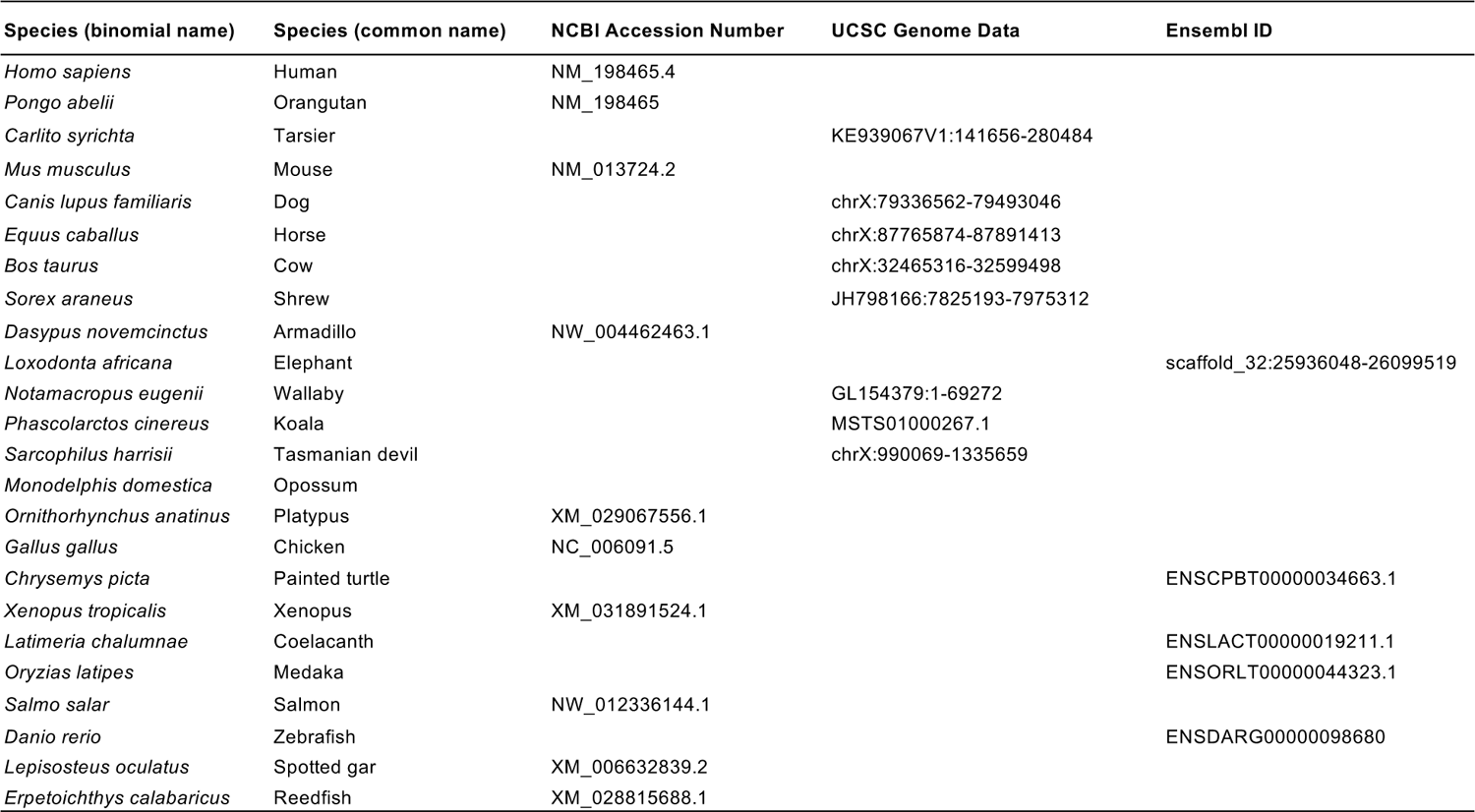
List of accession numbers of *Nrk* orthologs of selected species used in evolutionary analysis

**Supplementary Table 2.**
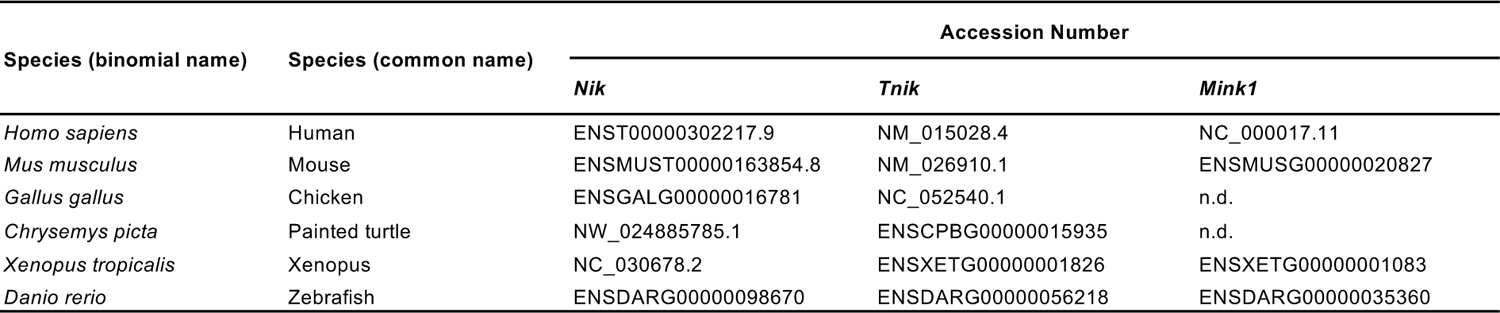
List of accession numbers of *Nik, Tnik, Mink1* of selected species used in synteny analysis

| Species (binomial name) | Species (common name) | Accession Number |  |  |
| --- | --- | --- | --- | --- |
|  |  | *Nik* | *Tnik* | *Mink1* |
| *Homo sapiens* | Human | ENST00000302217.9 | NM_015028.4 | NC_000017.11 |
| *Mus musculus* | Mouse | ENSMUST00000163854.8 | NM_026910.1 | ENSMUSG00000020827 |
| *Gallus gallus* | Chicken | ENSGALG00000016781 | NC_052540.1 | n.d. |
| *Chrysemys picta* | Painted turtle | NW_024885785.1 | ENSCPBG00000015935 | n.d. |
| *Xenopus tropicalis* | Xenopus | NC_030678.2 | ENSXETG00000001826 | ENSXETG00000001083 |
| *Danio rerio* | Zebrafish | ENSDARG00000098670 | ENSDARG00000056218 | ENSDARG00000035360 |

